# The effect of chronic stress and chronic alcohol intake on behavior, brain structure, and functional connectivity in a rat model

**DOI:** 10.1101/2025.02.13.638122

**Authors:** Jalil Rasgado-Toledo, Diego Angeles-Valdez, César J. Carranza-Aguilar, Alejandra Lopez-Castro, Luis A. Trujillo-Villarreal, David Medina-Sánchez, Mariana S. Serrano-Ramirez, A. Débora Elizarrarás-Herrera, Sarael Alcauter, Ilse Delint-Ramirez, Ranier Gutierrez, Gabriel A. Devenyi, M. Mallar Chakravarty, Eduardo A. Garza-Villarreal

## Abstract

Pathological chronic stress is stress exceeding the organism’s ability to cope physiologically, which may act as a risk factor in the onset and relapse of alcohol use disorder. Chronic- restraint stress (CRS) and ethanol intake are independently known to induce changes in brain structure and function, however, their combined effects on neurodevelopment over long periods of time remains largely unexplored. We conducted an in vivo longitudinal rat model with three main goals. 1) to determine if chronic stress increases ethanol intake; 2) to determine the effect of chronic- stress and ethanol intake in behavioral measures, brain structure, and function; and 3) to investigate the effect of sex. This observational study included Wistar rats assigned to four groups: 1) ethanol consumption (EtOH+/CRS-), 2) stress exposure (EtOH-/CRS+), 3) both ethanol and stress exposure (EtOH+/CRS+), and 4) control group (EtOH-/CRS-). Our results showed that chronic stress did not affect ethanol intake but led to reduced body weight gain, elevated corticosterone levels, and impaired recognition memory. Structural MRI revealed that both exposures produced additive brain volume changes in regions such as the olfactory bulb, orbitofrontal cortex, caudate-putamen, hippocampus, and cerebellum. Functional connectivity analysis using network-based statistics identified disrupted cortical-subcortical connections. Results found here were sex-dependent in terms of volumetric changes (higher effects on males) and functional connectivity (higher effects on females). Findings suggest sex-dependent mechanisms where both chronic- ethanol intake and stress affect brain plasticity during neurodevelopment. Understanding these region-specific vulnerabilities is crucial for addressing alcohol use disorders and stress-related neuropathology.

**Graphical abstract.**
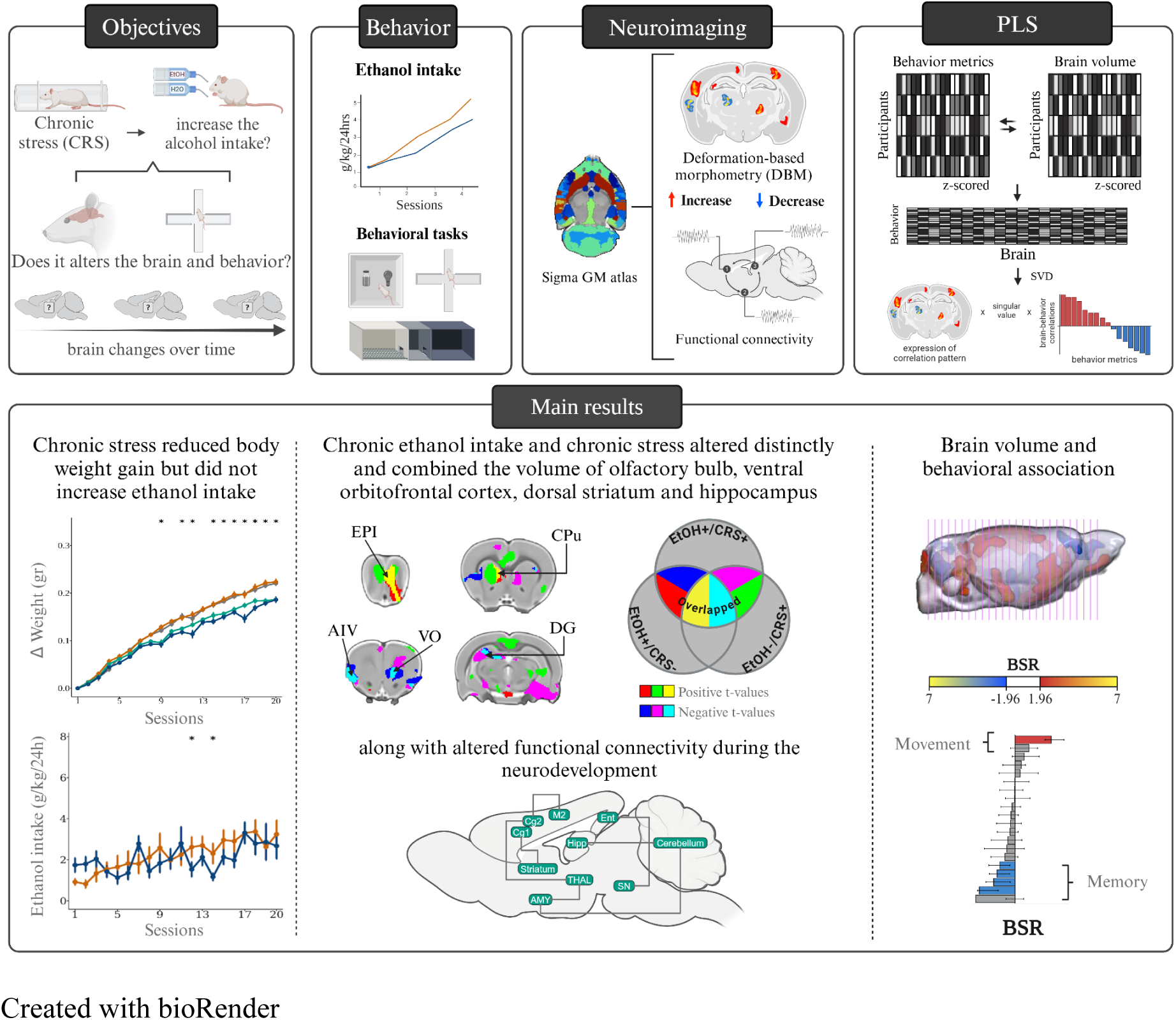
Created with bioRender

## 1. Introduction

Stress can be defined as the physical or psychological result of an organism’s exposure to adverse experiences and internal or external demands. Based on the chronology, it can also be defined as acute or chronic, the latter of which can become pathological.(1) Pathological chronic stress is described as stress exceeding the organism’s ability to cope physiologically, which may act as a risk factor in the onset and relapse of several neurological and psychiatric disorders, including substance use disorders (SUDs).(2–4) The chronicity of stressful events lead to a disruption of the finely regulated hypothalamic-pituitary-adrenal (HPA) axis and norepinephrine/autonomic systems (LC-NA), leading to neurochemical alterations associated with stress responses.(5,6) This disruption eventually entails an increase in behavior changes, such as alterations in anxiety levels, negative emotions, altered sleep patterns and appetite fluctuations, aggressive behaviors, and shifts in attention, concentration, and memory.(7) These disruptions often can lead to substance use and abuse as a way to cope with these negative emotional states.(8)

Alcohol use disorder (AUD) has been largely associated with acute and chronic stress, where it is commonly reported in humans that stress induces alcohol use, and that it may be used as a coping mechanism.(2,4) Several rodent studies have shown that exposure to acute and sub-chronic stress increased alcohol (ethanol) intake. However, other studies have shown the opposite, namely acute and sub-chronic stress exposure reduced ethanol intake. Furthermore, only a few studies have investigated the effects of chronic stress, mainly finding it has no effect on ethanol intake.(9) The possible mechanisms underlying stress and ethanol intake are not well defined and seem to be complex. Whether stress induces changes in ethanol intake seems to be dependent on the type of stressor, its intermittency, predictability, chronicity, and biological variables such as genetics, age, and sex.(2,9,10) Despite this complexity, the underlying and shared neural circuits could provide a unifying explanation for the relationship between stress and ethanol intake.(7) This has been suggested by neuroimaging studies in humans showing that changes in volume and function of specific brain regions can predict AUD development and relapse.(7,11–14)

Neuroimaging studies in rodent models of AUD have shown that chronic ethanol intake reduced the volume of specific brain regions such as hippocampus, thalamus, ventral tegmental area, substantia nigra, caudate-putamen, nucleus accumbens, retrosplenial, insular and prelimbic cortex, and increased the volume of orbitofrontal cortex, thalamus and cerebellum.(15–18) Functional neuroimaging (fMRI) studies revealed that chronic ethanol intake also induces a decrease in connectivity patterns between prefrontal cortical subregions, frontostriatal connectivity, and retrosplenial-visual networks, whereas between the prefrontal to cingulate and striatal networks increases.(19,20) In chronic stress, studies have shown it reduces volume in the hippocampus, prelimbic, cingulate, insular, somatosensory, motor, auditory and perirhinal/entorhinal cortices, striatum, nucleus accumbens, the bed nucleus of the stria terminalis, amygdala, and the thalamus.(21–24) fMRI studies have shown increased connectivity within functional networks such as the somatosensory, visual, and default mode(25), and increased connectivity of thalamic connections to the hippocampus, amygdala, ventral tegmental area, prelimbic, insular and retrosplenial cortices.(21) Hence, individual studies suggest brain regions that seem to be affected in both pathologies. However, there are no studies that investigate both at the same time and the possible synergy between them.

Sex-differences on the long-term influence of stress on ethanol intake have been described but are not well understood. For the most part, human research has described different coping mechanisms for SUD-related drinking behavior, possibly due to sexual differences in terms of neurochemical transmission, genetics, brain morphology, and connectivity.(26) New insights derived from different species addressing the ethanol influence in the context of intoxication, withdrawal, and cravings showed age and sex may act as mediators in different directions by either increasing or decreasing vulnerability to AUD and may serve as predictors of the negative impact on the brain.(27) Moreover, chronic stress studies have emphasized that the sex-specific impacts are also dependent on the drinking onset age, possibly driven by gonadal hormone fluctuations and the subsequent HPA/LC-NA responsiveness that occur over dynamic periods of development and maturation.(28)

Considering these considerations, our study goals were first, to determine if chronic stress increases ethanol intake; second, to determine the effect of chronic stress and chronic ethanol intake in behavioral measures, brain structure and connectivity; and third, to investigate the effect of sex. To this end, we employed a longitudinal design and *in vivo* magnetic resonance imaging in rat models of chronic stress and chronic ethanol intake.

## 2. Materials and methods

### 2.1. Animals

A total of 96 Wistar rats (45 females) were included in the experiment. The study was conducted over six batches of animals, with approximately 16 rats per batch. The rats were divided randomly into four experimental groups: Group 1 ethanol consumption (*EtOH+/CRS-*, n = 28, 13 females) consisted of animals under the Intermittent Access 2-Bottle Choice protocol (IA2BC, details can be found below) without the CRS; Group 2 stress exposure (*EtOH-/CRS+,* n = 20, 10 females) included animals that were not in the IA2BC but had the CRS protocol; Group 3 ethanol and stress exposure (*EtOH+/CRS+,* n = 28, 14 females) included animals under both the IA2BC and CRS protocols; and Group 4 control (*EtOH-/CRS-,* n = 20, 8 females) consisted of animals with no interventions (Figure 1a). For Group 4, two bottles were presented with only water to control for the IA2BC conditions.

**Figure 1.**
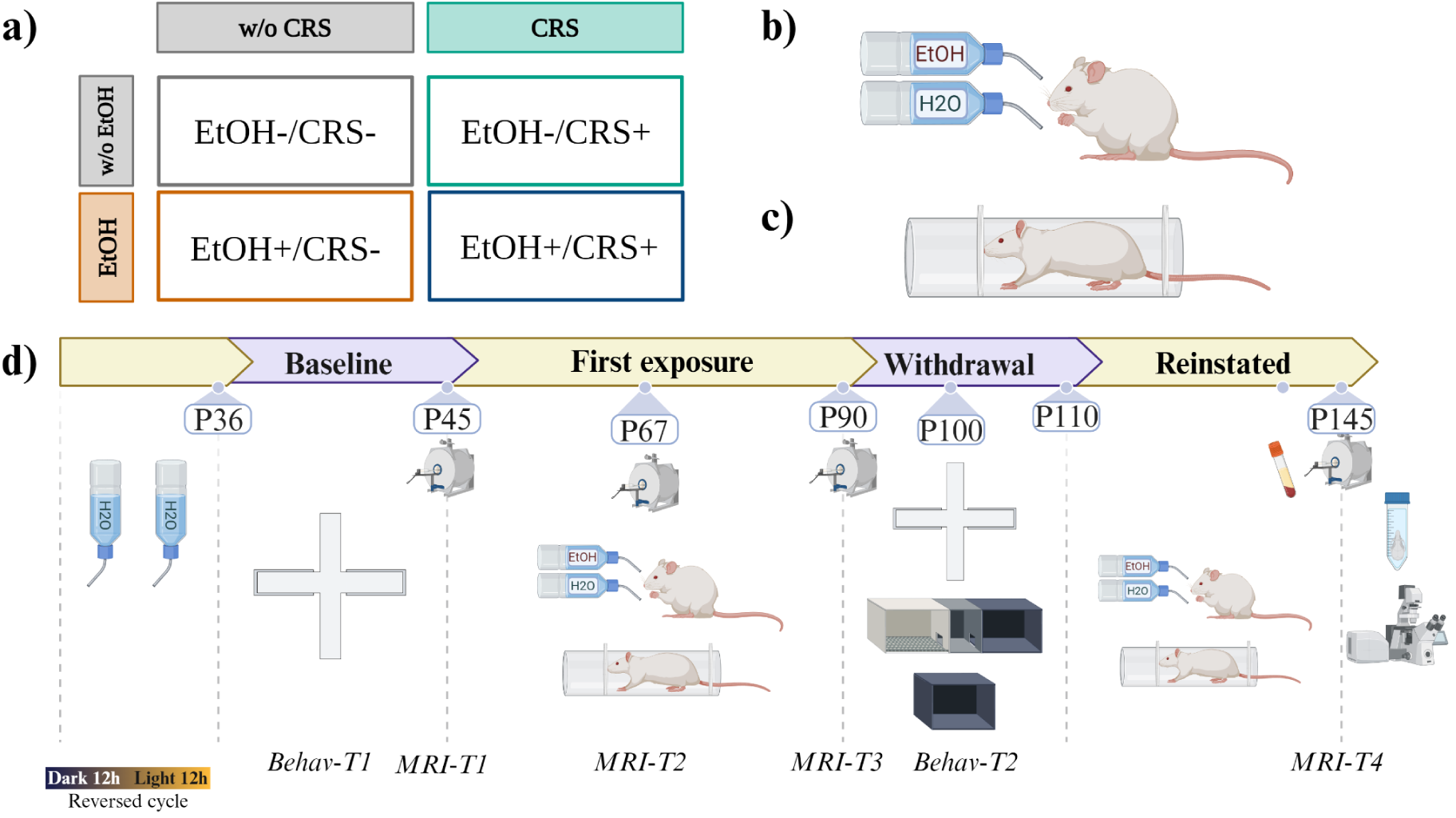
Experimental design. **a)** Representative group subdivision. **b)** Schematic drawing illustrating the intermittent access model. **c)** Schematic drawing of movement restrictor with adjustment for rat size. **d)** Experimental protocol: P21, rats were received; P35, baseline behavioral evaluation (Bevah-T1); P44, first MRI acquisition (MRI-T1); P45, initiation of IA2BC model during 45 days; P67, second MRI acquisition (MRI-T2); P91, third MRI acquisition (MRI-T3); P100, second behavioral evaluation (Bevah-T2); P91 to P110, withdrawal phase; P110, reinstaining of both interventions; P145, four MRI acquisition (MRI-T4). Abbreviations: ethanol intake (EtOH), chronic restraint stress (CRS), water intake (H2O), magnetic resonance imaging (MRI). Created with BioRender.

All experimental procedures and animal care were carried out in accordance with the “Reglamento de la Ley General de Salud en Materia de Investigación para la Salud” (Health General Law on Health Research Regulation) of the Mexican Health Ministry, which follows the “Guide for the care and use of laboratory animals”(29) and the “Norma Oficial Mexicana” (NOM-062-ZOO-1999). The animal research protocols performed were approved by the ethical committee of the Neurobiology Institute of the National Autonomous University of Mexico, project number A113. Appropriate procedures were performed to minimize animal suffering. This study adhered to the ARRIVE 2.0 guidelines for reporting animal research.(30)

### 2.2. Intermittent Access 2-Bottle Choice

The IA2BC protocol was adapted from Simms et al,.(31–33) (Figure 1b). Rats could consume water or ethanol (20%) on Monday-Wednesday-Friday. Twenty-four hours after placement, the EtOH bottle was weighted to obtain the binge (g/kg/30-min units), main intake (g/kg/24-hour units), and substance preference measures. Details about the description and procedures can be found in supplementary methods and in the workgroup’s repository(34).

### 2.3. Chronic restraint stress (CRS)

The chronic stress protocol was conducted using movement restraint as a stressor according to established protocols.(35) Movement restriction was performed intermittently for three hours from 11:00 (± 30 min) to 14:00 (± 30 min), five times per week. The stress days were selected randomly in order to diminish the habituation process.(36) The motion restrictor was constructed of acrylic with a double adjustment for the size of the rat (Figure 1c). All rats were weighed daily as a measure of stress response(35). Weight normalization was performed by subtracting the weight at P45 from all weight values according to age to obtain the weight change.

### 2.4. Blood corticosterone concentration

Blood samples were obtained from the rat tail at the end of the protocol (P142) in order to validate and verify CRS increasing corticosterone levels, as have been previously reported(37) (Figure 1). Details can be found in supplementary methods.

### 2.5. Experimental Design

The experimental design is outlined in Figure 1d. The protocol was divided into four phases: 1) Baseline, animals underwent initial behavioral testing (Behav-T1) at P36, followed by the first MRI acquisition (MRI-T1) at P44 ± 1 day. 2) In the first exposure phase, rats were randomly assigned to one of the four groups for 45 days (20 sessions of alcohol and/or 30 days of CRS). Additional MRI acquisitions were performed during this phase at P67 ± 1 day (MRI-T2) and at P91 ± 1 day (MRI-T3). 3) Withdrawal phase, absence of alcohol intake, and chronic restraint from P91 to P110; a second behavioral test was performed during this period, starting at P100. 4) In the reinstated phase, at P111, both interventions were implemented again. A final MRI acquisition was made at the end of the phase, at P145 ± 1 day (MRI-T4). Finally, animals were sacrificed (P146), and the brain was extracted for immunohistochemistry purposes, which were not analyzed here.

### 2.6. Behavioral tests

To measure the long-term outcomes of chronic stress and ethanol intake on anxiety-like behavior and recognition memory, we evaluated the animal’s behavior during the withdrawal phase using the elevated plus maze (EPM) and novel object recognition (NOR) tasks. We also evaluated the preference for alcohol over water using the place preference test (CPP). All behavioral tests were conducted under environmental conditions similar to those of the housing but with dim red lighting. Videos were preprocessed using ffmpeg to ensure consistent dimensions and object positions. DeepLabCut v. 2.3.8(38) was employed for automated behavioral tracking for EPM and NOR (Network training details can be found in Supplementary methods). Data tracking generated by DeepLabCut tracking was further processed using DLCAnalyzer(39) in R programming language (see Code Availability section for details). Detailed details of the statistical analysis can be found in the supplementary methods.

### 2.7. Magnetic Resonance Imaging analysis

#### 2.7.1. Structural analysis

All MRI scans were converted from Bruker format to NIfTI using the *brkraw* toolbox v0.3.11(40) using the BIDS framework(41). We then preprocessed the T2w images with intensity normalization(42), centering of the image, and denoise by using an in-house pipeline built on MINC-toolkit-v2 and ANTs tools and based on previous studies(43). One T2w image (1 subject-1 session) was excluded due to the high level of artifacts during the quality control stage. Finally, we performed Deformation Based Morphometry (DBM) employing a Two-Level DBM approach with the SIGMA anatomical template v. 1.2.1(44). We analyzed the voxel-wise longitudinal volume changes to find possible interactions between group, age, and sex. We used relative volume (Jacobian determinant relative to the averaged template created) per voxel as the dependent variable and age, group, and sex as independent variables.

Finally, to analyze the relationship between MRI brain volume and behavioral outcomes, we employed partial least squares (PLS) correlation (45). This multivariate approach allowed us to identify distributed patterns in brain structure and function related to behavior. For this analysis we associated the MRI-T3 with Behav-T2 acquisition points as they were acquired in adjacent time points.

#### 2.7.2. Functional analysis

For the resting state functional MRI (rsfMRI) data preprocessing, we employed the open-source RABIES pipeline for multi-stage correction.(46). To identify subnetworks (connected components) in rsfMRI that differ significantly by alcohol and stress conditions while controlling for multiple comparisons across a large number of connections, we used the R-library network-based R-statistics (NBR) that allows to perform network based statistics (NBS) with linear mixed-models.(47,48). We identified a connected subnetwork altered by the CRS (Figure 5a), and then, as a post hoc analysis, we focused on how the functional connectivity (FC) trajectory changed for each connection (ROI-to-ROI) within this subnetwork, filtering the most relevant connections by selecting the connections with significant group differences. Again, we used linear mixed-models with group, age, and sex interactions, batch as the covariate, and subject ID as a random effect (Eq. 1).

### 2.8. Statistical analysis

All statistical tests were performed through linear mixed-models or linear models in the case of novel object recognition. The local volume, functional correlation, weight change, ethanol intake, anxiety index, and preference index were considered as dependent variables to test group differences (Eq. 5, see supplementary methods). Group, session, and sex were included as interaction fixed effects (independent variables) and rat identification (RID) as a random effect. Batch was used as a covariate in all models. In contrast, the discrimination ratio was analyzed using a linear regression model (Eq. 6, see supplementary methods).

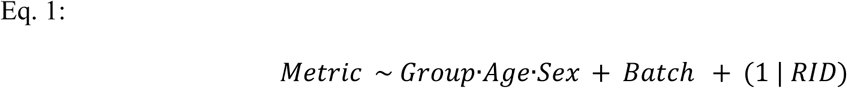

Metrics: local volume, functional connectivity, ethanol main intake, Δweight = (weight in each session - weight of the first session), anxiety index, and preference index.

For all analyses, alpha level was set at 5%, and for q = 0.05, false-discovery (FDR). All scripts to reproduce the analysis and ROI-specific masks are available in the GitHub repository (See code availability section). Detailed details of the statistical analysis can be found in the supplementary methods.

## 3. Results

### 3.1. Ethanol intake

The significant pairwise comparisons of the IA2BC model indicated specific group differences in the main intake of ethanol across only some sessions (Figures 2a and 2c). Chronic restraint stress (EtOH+/CRS+) reduced the ethanol intake compared to EtOH+/CRS- group in males after a total of 31 sessions (20 sessions of first phase and 11 of the reinstated phase), the preference of it over water was also diminished. The preference index by CPP task also showed a lower ethanol preference in male EtOH+/CRS+ group (Figure 2b and 2d). Interestingly, we also found that after the withdrawal period, the male ethanol group (EtOH+/CRS-) consumed larger quantities than before (rebound effect), which was not found in the EtOH+/CRS+ male group or both female groups.

**Figure 2.**
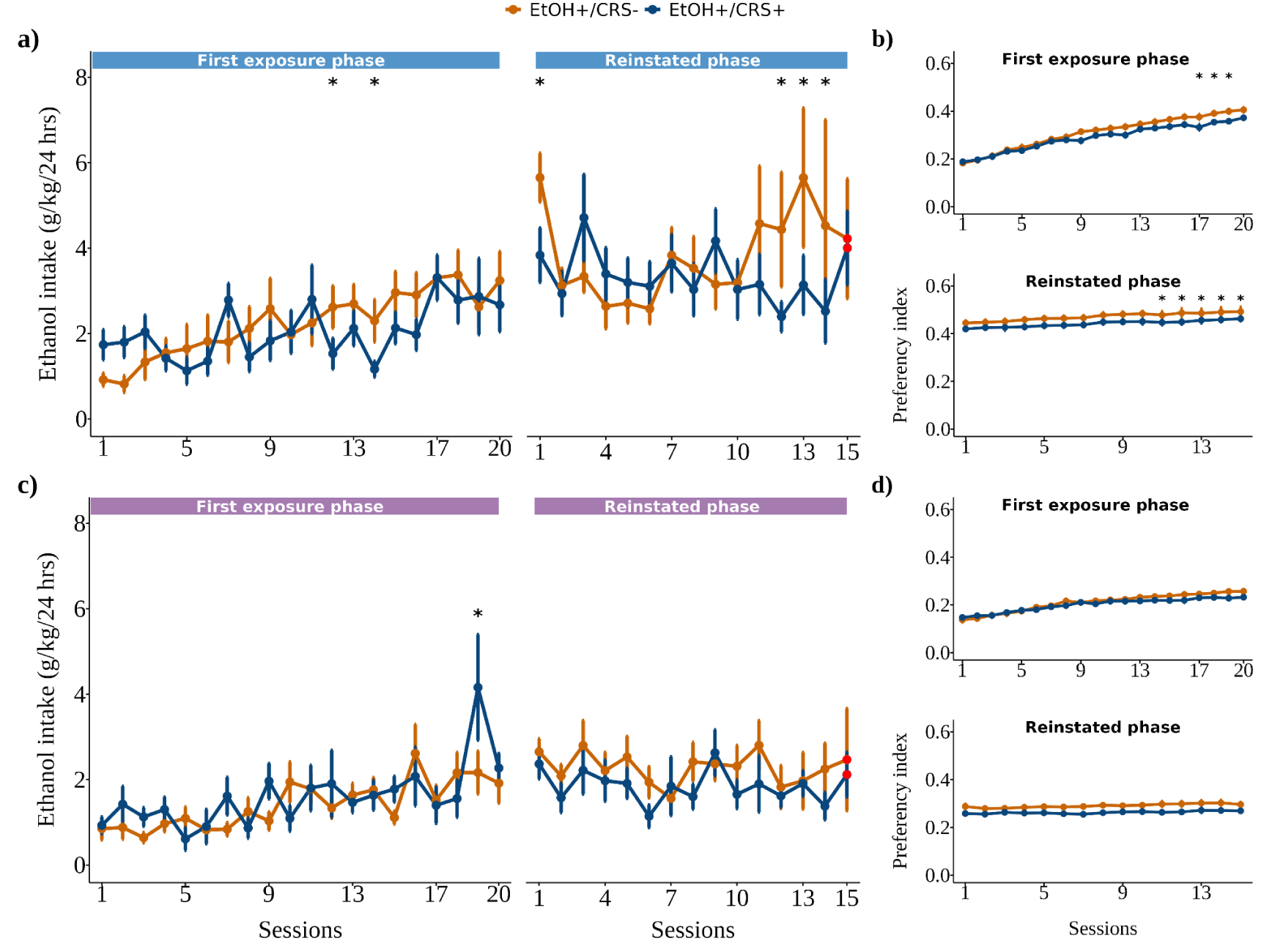
Ethanol intake and preference. Chronic stress reduces ethanol intake and CPP in males. **a)** Male and **c)** female ethanol intake over the twenty sessions of the first exposure and in the fifteen sessions of the reinstated phase, on the right, a boxplot representation of the median of all sessions. **b)** Male and **d)** female ethanol preference index of ethanol over water, the threshold above 0.5 means EtOH preference index. Group differences were assessed by a linear mixed model (See eq. 5, supplementary methods). Abbreviations: ethanol intake (EtOH), chronic restraint stress (CRS).

### 3.2. Weight

Throughout the sessions, the weight of the animals under CRS (EtOH-/CRS+, EtOH+/CRS+) reduced the body weight gain on both sexes compared to the control and ethanol intake group, even during relapse (Figures 3a and 3b).

**Figure 3.**
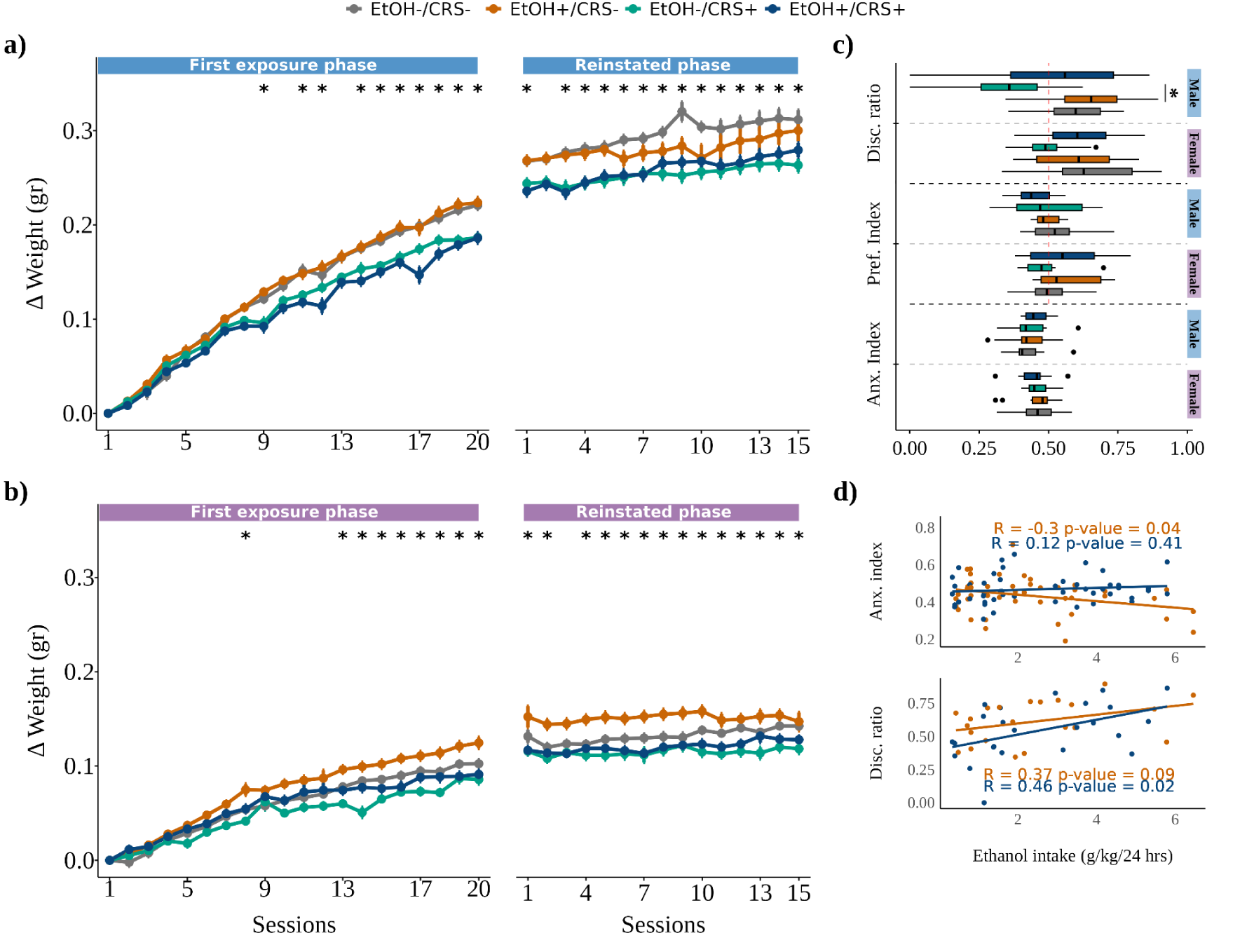
Body weight gain and behavior. Chronic stress reduces body weight gain and affects object recognition memory in males. **a)** male and **b)** female weight change over the twenty sessions of the first exposure and in the fifteen sessions of the reinstated phase. **c)** Indexes from the first behavioral evaluation (Behav-T1). **d)** Indexes from the second behavioral evaluation (Behav-T2). Anxiety index = calculated based on elevated plus maze task (Eq. 2). Disc. ratio = Discrimination ratio calculated based on novel object recognition task (Eq. 3), threshold above 0.5 means novel object interaction. Pref. Index = Preference index calculated based on conditioned place preference task (Eq. 4), threshold above 0.5 means ethanol chamber preference. Group differences were assessed by linear mixed model (See eq. 1).

### 3.3. Blood corticosterone concentration

As expected, corticosterone levels in blood serum were significantly higher in both stress groups compared with the control (EtOH-/CRS+: q = 0.006; EtOH+/CRS+: q = 0.004) and with EtOH+/CRS- group (EtOH-/CRS+: q = 0.013; EtOH+/CRS+: q = 0.006), which confirms the success of the stress protocol. Ethanol exposure did not influence corticosterone levels (See supplementary figure 4).

### 3.4. Behavioral test

The anxiety index obtained by the EPM task did not show any differential performance depending on the treatment (q > 0.05) or sex (q > 0.05), meaning none of the groups showed overall anxiety-like behaviors after chronic stress/chronic ethanol intake (Figure 3d). However, individual EPM metrics (See supplementary results) showed that females (EtOH+/CRS- and EtOH+/CRS+) who were exposed to chronic ethanol intake showed behaviors associated with anxiety, such as more stationary time during the task and less speed reached than the female’s control group, although the anxiety index (global measure) was not different between groups and sexes.

Discrimination ratio of the NOR task (Eq. 3) showed a lower novel object recognition only in the male EtOH-/CRS+ group compared with male EtOH+/CRS- (β = 0.26, q = 0.023, p = 0.003, d = 1.53), and uncorrected significant compared with male EtOH-/CRS- (β = 0.19, q = 0.096, p = 0.048, d = 1.1) and with male EtOH+/CRS+ (β = −0.17, q = 0.096, p = 0.048, d = −0.97) (Figure 3d). In contrast, individual measures (See supplementary results) showed that male and female EtOH+/CRS+ had lower exploration metrics of the novel object (more distance covered, reaching higher speed and moving time) than EtOH+/CRS- group. In summary, males with chronic stress exposure showed globally lower novel object recognition compared to the males with chronic ethanol intake exposure and male controls (See supplementary results).

### 3.5. Voxelwise analysis

The chronic ethanol intake and chronic stress (EtOH+/CRS+ group) showed an additive effect in cortical and subcortical brain regions such as olfactory bulb (OB), frontal association area (FrA), secondary motor area (M2), secondary cingulate cortex (Cg2), entorhinal cortex (Ent), cerebellum (Cer), caudate-putamen (CPu), hippocampus and thalamic areas (Figure 4a). This group (EtOH+/CRS+), compared with the ethanol intake (EtOH+/CRS-) group, displayed volume reductions in the frontal associative cortex (FrA), insular cortex (AIV), ventral orbitofrontal cortex (VO), retrosplenial cortex (RSGc), hippocampus. The increased volumes were found in the olfactory bulb, CPu, Ent, and cerebellum (Figure 4b). On the other hand, when we contrasted with the EtOH-/CRS+ group, we found a decrease in FrA, VO, CPu, hippocampus, hypothalamus, and amygdala (Amy), while increasing volumes were found in OB, CPu, Cg2, RSGc, Ent, cerebellum and thalamic areas (Figure 4b).

**Figure 4.**
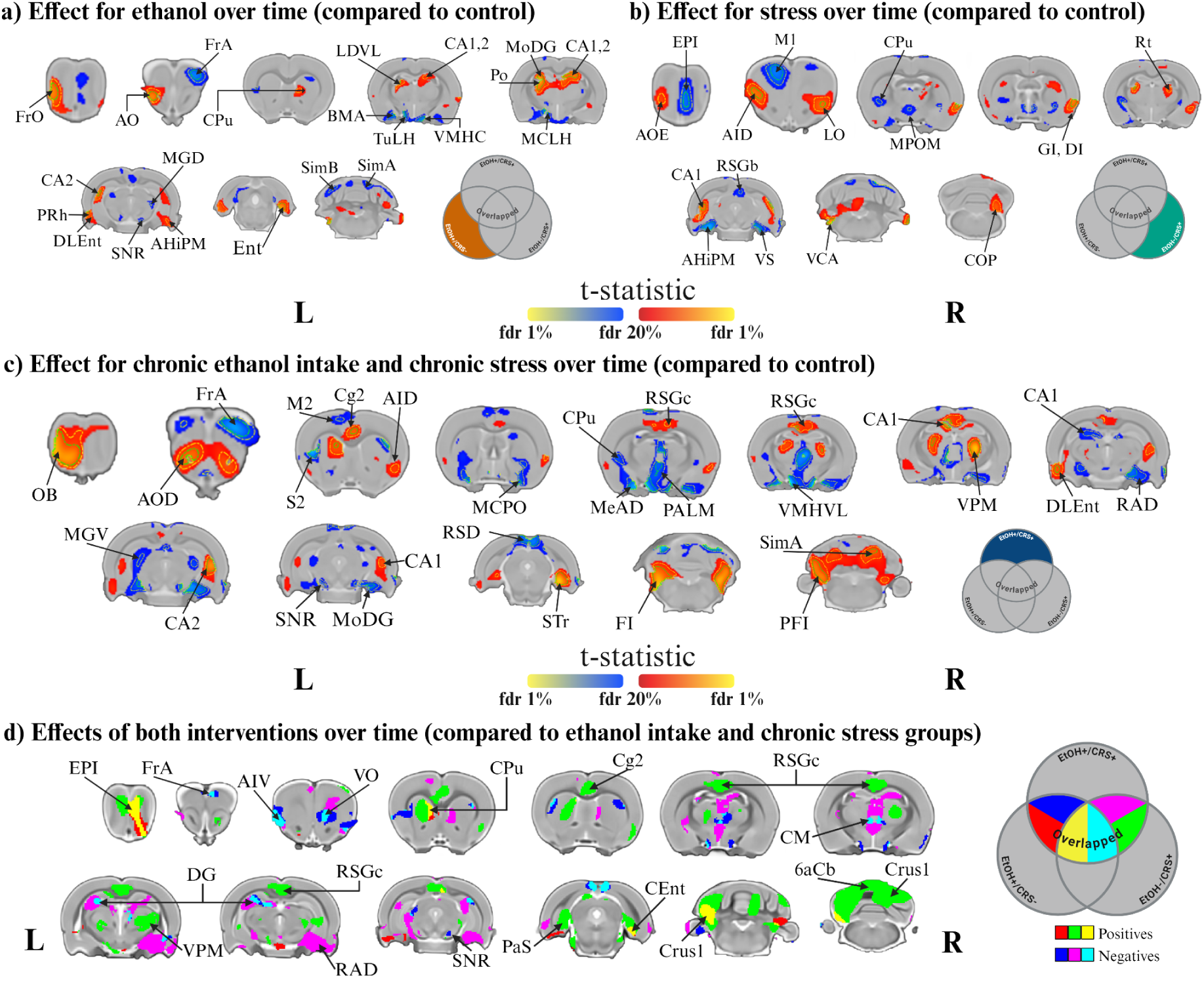
Structural effects of chronic ethanol intake, chronic restraint stress and both interventions among the control group (Interaction Age*Group*Sex). Chronic ethanol intake and chronic stress affected more extended areas contrasted with all groups. The effect appear to be additive (yellow and light blue) in olfactory bulb (EPI), ventral orbitofrontal cortex (VO), dorsal caudate-putamen (CPu) and dentate gyrus (DG) and cerebellum (Crus 1). **a)** Longitudinal structural effects of chronic ethanol intake (EtOH+/CRS-), **b)** chronic stress (EtOH-/CRS+) and **c)** the effects of both interventions (EtOH+/CRS+) over the control (EtOH-/CRS-). To explore the additive effect of each intervention in the EtOH+/CRS+ group, we contrasted this group against EtOH+/CRS- or EtOH-/CRS+. **d)** Additive effect of chronic ethanol intake (EtOH+/CRS+ > EtOH-/CRS+; red: positive t-values, blue: negative t-values), chronic stress (EtOH+/CRS+ > EtOH+/CRS-; green: positive t-values, purple: negative t-values) and overlapping effects of both. Color map describes the direction of the t-statistics; cooler colors denoting negative values; most commonly corresponding to volume decline and warmer colors denoting positive values. Threshold set at FDR of 20%, 5% (yellow dashed line) and 1% (green dashed line). Coronal slides labels according to Paxinos & Watson stereotaxic atlas (62).

Overall, we found that the male rats that were exposed to chronic stress and chronic ethanol intake showed higher extension of longitudinal volume changes compared to all groups and females. The effects appear to be additive in the olfactory bulb (EPI), ventral orbitofrontal cortex (VO), dorsal caudate-putamen (CPu), dentate gyrus (DG), and cerebellum (Crus 1) when compared with the groups with only one intervention and females. Males, in general, are more affected than females in terms of local volume.

### 3.6. Structural brain volume and behavior relationship

Partial least squares results showed one significant latent variable (LV-3) that explained 17% of the variance (p = 0.011) with significant brain correlation (See supplementary results), while the rest of the components explained most of the variance but were not significant results (Figure 5a). The post-hoc analysis of the PLS did not show group differences in the behavior scores nor the brain scores (Figure 5b,c), therefore the association results do not refer to one group in particular. We found the brain-behavior metrics association between the moving time (percentage) in the familiar object (NOR) with the volume of the prelimbic, orbitofrontal, insular, motor, primary cingulate, somatosensory, visual cortices along with the ventral striatum (CPu) and anterior regions of the cerebellum. While total time spent (NOR), the stationary time with the novel object, and the total time and transitions to the familiar object were associated with the olfactory bulb, frontal associative and somatory cortices, striatum, hypothalamus, thalamus and posterior cerebellum subregions (Figure 5d,e). In summary, the PLS-analysis found a significant association between locomotor activity with widespread cortical regions and recognition memory-related metrics with mostly subcortical and cerebellum regions in all groups (See supplementary results).

**Figure 5.**
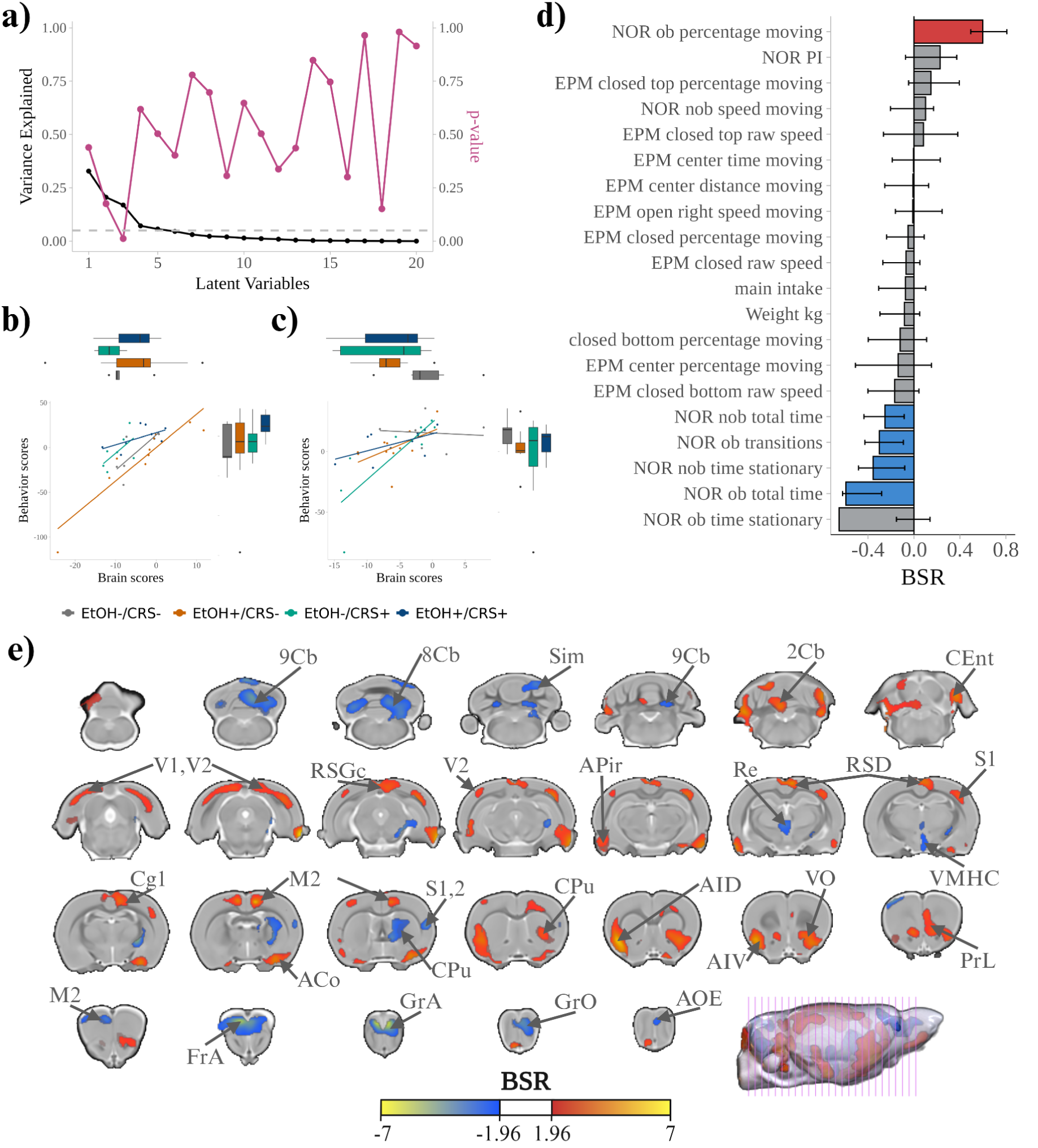
Relationship between the brain volume and EPM-NOR-behavior of the latent variable 3 (LV3). **a)** The third LV explained 17% of the covariance between brain and behavior metrics (p = 0.011). Dashed line at 0.05. Pairwise group comparisons of the PLS brain and behavior scores of LV3 on **b)** males and **c)** females. *q < 0.05, **q < 0.01, ***q < 0.001. **d)** Spatial brain patterns of the contribution to the bootstrap ratio (BSR) values. Cold colors indicate negative BSR values, while warm indicates positives (threshold set to 1.96). **e)** Behavioral metrics patterns associated with the brain pattern. Bars are colored to show significant association. The brain pattern is associated with moving-related metrics in positive BSR values, while negatives are associated with memory-related metrics.

### 3.7. Functional connectivity

The NBS linear mixed-effects analysis (Eq. 7) revealed significant effects in the EtOH-/CRS+ group compared to the EtOH-/CRS- group, identifying 46 connections across 37 regions (q =0.032, Figure 6a). Subsequent post-hoc testing of significant group differences in functional connectivity (FC) identified seven altered edges involving the amygdala, thalamus, caudate-putamen, substantia nigra, hippocampus, cerebellum, entorhinal cortex, secondary motor cortex, and primary and secondary cingulate cortex (Figure 6b-h).

**Figure 6.**
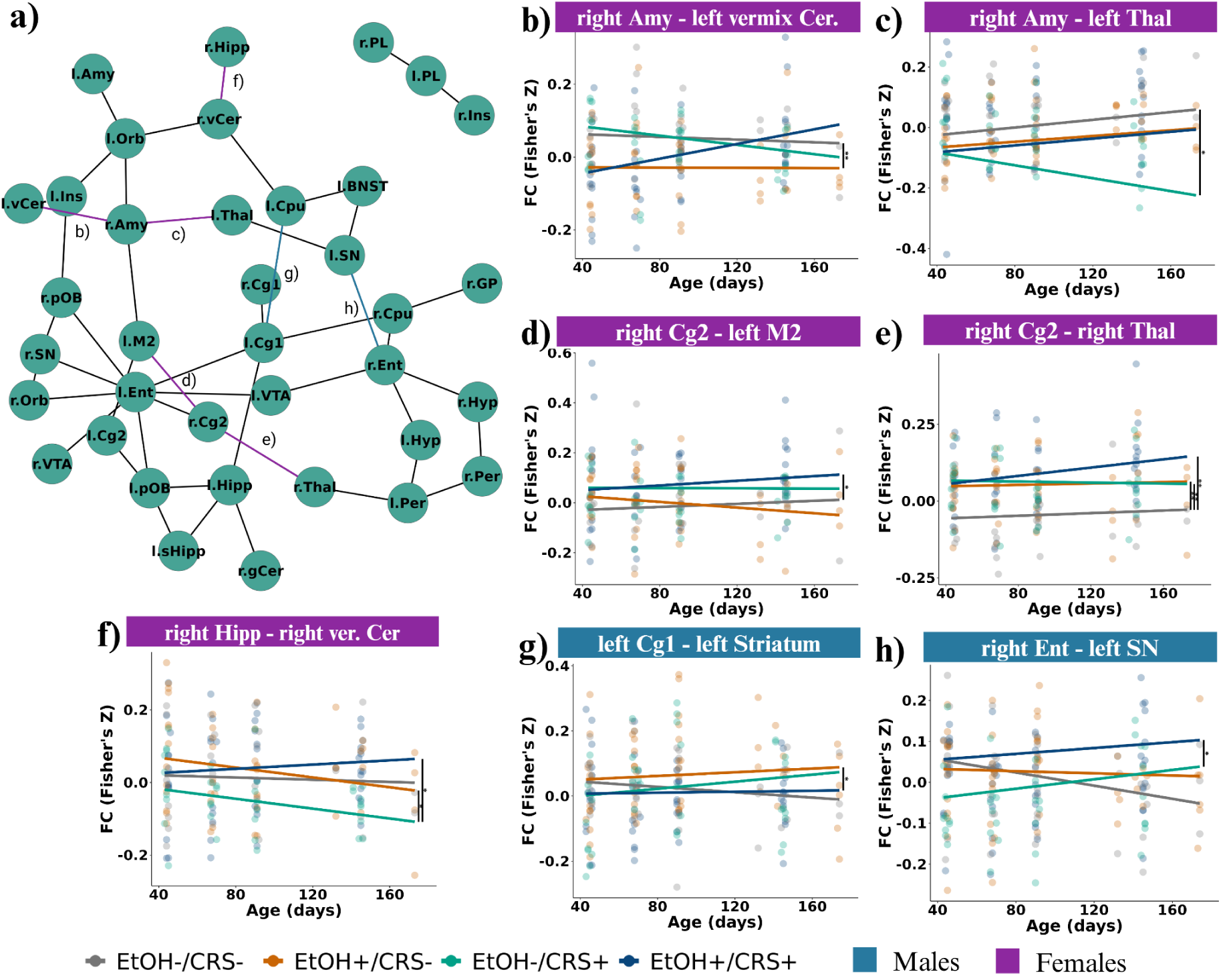
Functional connectivity effects of chronic ethanol intake and chronic restraint stress (EtOH+/CRS+) contrasted with the other groups in male rats (Interaction Age*Group*Sex). Chronic stress altered the functional connectivity of 37 regions compared with the control group, while both interventions (EtOH+/CRS+) changed the FC of more connections over the neurodevelopment. **a)** NBR analysis via linear mixed model analysis (Eq. 7) results in regions (nodes) comprising the brain network with significant changes in functional connectivity (Fisher z-transformed correlation values) along the age. **b-h)** Scatter plot (residuals) with regression lines showing the trajectory of each connection. Abbreviations: secondary cingulate cortex (Cg2), secondary motor cortex (M2), primary cingulate cortex (Cg1), entorhinal cortex (Ent), hippocampus (Hipp), thalamus (Thal), amygdala area (Amy), substantia nigra (SN).

Compared to the control group, we found distinct FC patterns in all experimental groups. Chronic ethanol intake alone altered the FC of only one element (right amygdala - cerebellum), while chronic stress alone and combined with ethanol intake altered FC in more connections compared to the control and between these groups. The female EtOH+/CRS- rats exhibited altered connectivity between the right amygdala and the left vermis of the cerebellum (right Amy-left vCer, figure 6b), while the female EtOH-/CRS+ group showed altered connectivity between the right amygdala and the left thalamus (right Amy-left Thal, figure 6c). The female EtOH+/CRS+ group showed altered significant connectivity vs the control in the connection of the right secondary cingulate cortex with the left secondary motor cortex (right Cg2-left M2, figure 6d). The three experimental groups displayed altered connectivity between the right secondary cingulate cortex and the right thalamus (right Cg2-right Thal, figure 6e), but only the EtOH+/CRS+ group had altered connectivity of right Cg2 with the left secondary motor cortex. Additionally, both female ethanol groups showed altered connectivity between the right hippocampus and the right vermis of the cerebellum (right Hipp-right vCer, figure 6f) contrasted with EtOH-/CRS+. On the other hand, only male EtOH+/CRS+ rats exhibited altered connectivity between the left primary cingulate cortex and the left striatum (left Cg1-left Str, figure 6g) in comparison to the male EtOH+/CRS- group, as well as between the right entorhinal cortex and the left substantia nigra (right Ent-left SN, figure 6h) in contrast to the male EtOH-/CRS+ group. Overall, we found that chronic stress altered the functional connectivity on 37 cortical and subcortical regions compared with the control group. Increased connectivity was found by the combination of both interventions mostly in cortical to subcortical connections (Figure 6d-h, supplementary table 1). This effect was present in females more than males, which is opposite to the structural results.

## 4. Discussion

In this study, we aimed to determine if chronic stress increases ethanol intake, to understand the effects of chronic stress and chronic ethanol intake on behavioral tasks, brain structure, and function, and to investigate the effect of sex. Our study demonstrated that chronic stress had no significant impact on ethanol intake for the initial 20 sessions of the first phase, only two days were different but not consecutive (see Figure 1a). However, beyond 11 sessions during the reinstated phase (31 sessions in total), chronic stress was associated with a reduction in ethanol intake. We also found that chronic stress reduced body weight gain, even in the chronic stress/ethanol intake group, increased corticosterone levels and decreased the recognition memory index. Neuroimaging analysis showed that the combination of chronic stress and ethanol intake induced the most prominent neuroanatomical changes in the brain, with apparently additive effects in the olfactory bulb, caudate-putamen, orbitofrontal, hippocampus, and cerebellum, more affected on males. Opposite to this result, we found functional connectivity changes in several networks, such as amygdala-thalamus and cingulate-striatum, hippocampus-cerebellum, among others, more affected on females. Overall, our results show a complex relationship between chronic stress, ethanol intake, and sex that should be studied further.

Contrary to some animal studies, we did not find that chronic stress increases ethanol intake(9), but we found a tendency for a decrease in ethanol intake after 31 sessions. Previous works have discussed the contradictory results in acute and subchronic stress studies. Some studies have found that stress increases ethanol intake, others have found that stress reduces ethanol intake, and others have found no effect.(9) In our study, we found that chronic stress reduced ethanol intake in the last three sessions only in males, suggesting a possible dynamic relationship between both conditions that cannot be studied in short time frames and that chronic stress may play a role in ethanol intake in the latter stages. There may also be an effect of onset age of chronic stress and/or ethanol intake, where the onset of either condition in adolescence shows a different dynamic than in adulthood. Our study is also the first one to investigate chronic stress and chronic ethanol intake together from adolescence to adulthood. Therefore, our results are novel and should be further replicated, specifically testing the onset age of either or both conditions. We also found that males consumed more ethanol than females but only in the reinstated phase,(49,50) supporting the idea that sex is related to the vulnerability to AUD but is also age onset and abstinence-dependent. (27)

Voxel-wise local volume analysis revealed region-specific changes in response to chronic ethanol intake, chronic stress, or their combination, highlighting distinct neurodevelopmental trajectories in rats that seem to be sex and age onset-dependent. The overlapping volume changes were found in the olfactory bulb (OB), ventral orbitofrontal cortex (VO), caudate-putamen (CPu), dorsal hippocampus (Hipp), and cerebellum (Cer), previously found in human studies as overlapping regions influenced by ethanol intake or stress (8). Some authors have hypothesized that there may be additive effects of chronic stress and ethanol intake due to microstructural changes such as cell degeneration.(51,52) However, this has not been directly studied in animal models. Moreover, our functional connectivity (FC) results showed a disrupted trajectory pattern between the connectivity of cortical to subcortical regions such as cingulate to striatum, -thalamus, and -secondary motor cortex in the combined chronic stress and ethanol intake in females, but more so in the chronic stress group. Chronic stress may produce an altered network integration across multiple brain regions, as some animal MRI studies have shown.(20,21) It seems that the more robust structural changes that we see in male rats with both interventions may be the results of an additive effect, while the functional connectivity changes in females with chronic stress may be the results of neuroadaptations related to this particular intervention. The additive effect of both interventions over core regions implicated in addiction and stress-related circuitry could serve as potential target regions to understand the intricate mechanisms of stress effects over ethanol intake. Moreover, the behavioral results showed a different pattern in the locomotor activity and recognition memory, more affected by chronic stress (further discussion can be found in Supplementary files). Interestingly, the association between volume and behavior displayed two metric sub-grouping. Locomotor activity metric associated with the volume of cortex and cerebellum regions, typically associated with this process (53–55). While olfactory bulb, thalamus, dorsal striatum were found to have associated with recognition memory-related metrics which were found altered by the chronic stress (55,56). However, contrary to our hypothesis, the brain-behavior association was not specific to any group.

The study of sex differences in psychiatry has gained significant importance over the past five years, as much of the existing literature in animal models has excluded female subjects, often assuming that findings in males can be generalized to both sexes.(57) However, the latest studies suggest that there are clear sex differences that need to be addressed. For this reason, we included females in our study to address this complex issue. In psychiatry research, sex effects have been suggested to be related to hormones. Females have a different pattern of functionality in the HPA-axis, LC-NA system due to gonadal hormone fluctuations that lead to a different stress responsiveness.(28,58) Studies have observed that females have pulses or oscillations of stress-interacted hormones that are more well defined than in males and might have special relevance in stress-induced ethanol intake interaction.(59) Our results suggest that brain structure was less affected on females, but functional connectivity showed the opposite, being females the most affected. For this reason, chronic stress may differentially affect males and females in their behavior, brain structure and functionality.(28,60) This assumption needs to be further studied using microstructural and other functional techniques such as exploring the neuronal function and inflammatory signaling pathways.(61) These findings should be interpreted with caution, given the limitations of animal models for studying disorders. In this study, we used only movement restraint to model the complex phenomenon of stress-induced ethanol intake, which may not fully capture the intricacies of this behavior. Moreover, although the neuroimaging data were collected longitudinally, our behavior data was cross-sectional and during an abstinence period. Future research should aim to extend these analyses across the lifespan.

## 5. Conclusion

Overall, the present results reflect neuroadaptive or maladaptive neuroplastic responses especially with prolonged restraint stress, disrupting cognitive behavior and altering the brain at macrostructural level, without any corresponding changes in ethanol intake levels. These results underscore the sex-dependent and region-specific effects of combined ethanol exposure and chronic stress on both the structural and functional networks of the brain, emphasizing the need for further studies to understand the mechanisms driving these interactions in the course of psychopathological development.

## 6. Data availability

The data that support the findings of this study are available on request from the corresponding author, E.A.G.V.

## 7. Code availability

For the code analysis and extended results presented here, please check: https://github.com/JalilRT/Sudmex_alcohol_stress

## Supporting information

Supplementary results

Supplementary discussion

## Acknowledgements

We appreciate the support of the vivarium of the Instituto de Neurobiología of the UNAM, staff: MVZ. José Martín García Servín, Dra. Alejandra Castilla León and Dra. María A. Carbajo Mata. Behavioral Analysis Department, staff: Dra. Deisy Gasca Martínez. Proteogenomics Department, staff: M. S. Adriana González Gallardo. Microscopy unit: Dra. Elsa Nydia Hernández and Dra. Ericka de los Ríos. MS. Soledad Mendoza Trejo for the value help on the experimental technical support. Juan Pablo Maya Arteaga and Diego Emmanuel Ortuzar Martínez for the help on data collection. We also appreciate the technical support of Dr. Juan Órtiz and Dr. Luis Concha from the National Laboratory of Magnetic Resonance Imaging (LANIREM). We also thank Leopoldo González, Luis Aguilar, and the technical staff from the laboratory Lavis (UNAM Juriquilla), and Computational department staff: Ing. Ramón Martínez Olvera and M. Moises Mendoza Baltzar. This research was enabled in part by support provided by M. Mallar Chakravarty and Gabriel A. Devenyi at the Computational Brain Anatomy Lab (CoBrA Lab) (http://cobralab.ca/), CIC, Douglas Research Centre, Montreal and Digital Research Alliance Canada (www.computecanada.ca) and Mallar Chakravarty (Director of the Computational Neuroanatomy Laboratory, Douglas Research Centre, Montreal, Canada) who provided access to computational tools from his group on the Niagara Compute Cluster.

Work funded by: Instituto de Neurobiología UNAM and project PAPIIT-DGAPA IA202120 and IA201622. Jalil Rasgado-Toledo is a doctoral student from the Programa de Doctorado en Ciencias Biomédicas, Universidad Nacional Autónoma de México (UNAM) and has received CONAHCYT fellowship number 858667.

