## Supplementary methods for "The effect of chronic stress and chronic alcohol intake on behavior, brain structure, and functional connectivity in a rat model"

### Animals

For habituation to the experimenters and the manipulation, every day for ten minutes the offspring rats were handled to control the novelty bias. On postnatal day 25, the pups were housed individually and provided with two bottles with 100 ml of water to get them accustomed to the bottles for the ethanol intermittent access 2-bottle choice (IA2BC) protocol.

Postnatal day 21 weaned Wistar rats (Rattus norvegicus albinus) were bred in the Institute of Neurobiology's vivarium in groups of four in polypropylene transparent cages. Rats were housed in a temperature (26°C), humidity (60%), and wood chip bedding of 2 cm in depth in the same-sex room under a 12-h dark/12-h light inverse cycle (light on between 19:00 and 07:00). Rat chow (LabDiet, Richmond, Indiana) and water were ad libitum.

### Intermittent Access 2-Bottle Choice

This model has shown that repeated cycles of ethanol intake and withdrawal under a free-choice design lead to a gradual escalation in intake, and has also been replicated with high validity and reliable results. Starting on postnatal day 46, rats began the IA2BC protocol (Figure 1b). Rats could consume water or ethanol (20%) on Monday-Wednesday-Friday. Twenty-four hours after placement, the EtOH bottle was weighted to obtain the binge (g/kg/30-min units), main intake (g/kg/24-hour units), and substance preference measures. Details about the description and procedures can be found in supplementary methods and in the workgroup's repository(35). All bottles used in this study had anti-drip nozzles (RedKit, 125 mL), and after every refill, a square of parafilm over the bottle ring was placed to reduce fluid loss.

Starting on postnatal day 46, rats began the IA2BC protocol. On Monday-Wednesday-Friday one 100 ml bottle contained (EtOH) 20% ethanol concentration in potable water, while the other 100 ml bottle held plain potable water ($H_2O$) (Figure 1b). On Tuesday-Thursday-Saturday-Sunday the two bottles had 100 ml of potable water. These days could be considered as withdrawal periods. The measures of ethanol intake were obtained on Tuesday-Thursday-Saturday in g/kg/24-hour units. The placement of the ethanol bottle was randomized in each drinking session to control for side preferences. Thirty minutes and twenty-four hours after placement, the EtOH bottle was weighted in order to obtain the binge, main intake, and substance preference measures (Eq. 1.):

Eq. 1:

$$Alcohol\, main\, intake = (Initial - 24hrs) \cdot Animal\, weight\ (\mathrm{g})$$

$$Binge = (Initial - 30min) \cdot Animal\, weight\ (\mathrm{g})$$

$$Preferency = Alcohol\, main\, intake \div Total\, fluid\, intake$$

### Blood corticosterone concentration

For this procedure, rats were immobilized using movement restriction tubes during the extraction process, which lasted no longer than 5 minutes to prevent any acute stress response caused by handling [1,2]. Once the blood samples were collected, they were placed on ice and immediately centrifuged at 5000 rpm at 4 C for 20 min (Eppendorf centrifuge, Model 5415R). To analyze serum corticosterone concentrations, we followed the instructions provided by the commercial enzyme-linked immunosorbent assay (ELISA) kit for corticosterone (ALPCO, Cat. 55-CORMS-E01). Using a spectrophotometer (Varioskan LUX Multimode Microplate Reader v.4.00; Thermo Fisher Scientific, MA, USA) absorbance was measured at 450 nm, and serum concentration (ng/ml) was determined through the four-parameter logistic equation (4-PL).

### Behavioral tests

Animals were moved an hour previous to each evaluation for habituation to room conditions. To avoid circadian variations, all experiments took place during the rats' dark/active cycle at the Institute's Behavioral Analysis Unit.

The EPM test apparatus consisted of a plus-shaped maze with two open arms (45 cm × 5 cm each) and two enclosed arms (45 cm × 5 cm each), connected by a central platform (5 cm × 5 cm). The maze was elevated 40 cm above the floor. Individual rats were evaluated and recorded for five minutes (HD camera model S4612). Each video was compressed and cropped to the dimensions of the maze for subsequent software analysis by using ffmpeg v.4.2.7.

DeepLabCut training network for EPM was created using 20 distinct frames randomly selected from each of 14 videos of different animals. Each frame was manually labeled in a top-down view for each segment of the maze (top, down, left, right) and eleven body parts of the rats (nose, head center, neck, body center, body center left, hip joint left, hip joint right, tail base, tail center, and tail tip). New videos were continually added, requiring re-labeling and re-training of the network. A ResNet-50 based neural network with default parameters was used for training with a total of 2,030,000 iterations. The trained network achieved a test error of 2.17 pixels and a training error of 2.64 pixels. With a p-cutoff value of 0.6, the training error improved to 2.45 pixels, and the test error decreased to 1.9 pixels. Data tracking generated by DeepLabCut was further processed using DLCAnalyzer [3] in R programming language (see Code Availability section for details). The distance traveled, velocity, number of entries into each zone (open arms, closed arms, and center platform), and time spent in each zone were based on the data tracking. The time spent and number of entries in each arm were used to calculate an anxiety index (Eq. 2). A higher anxiety index value corresponds to greater anxiety-like behavior in the rat.

Eq. 2:

$$Anxiety\ index\ = 1\ -\ \left(\left[\frac{Open\ arms\ time}{Total\ time}\right]\ +\ \left[\frac{Open\ arm\ entries}{Total\ entries}\right] \div 2\right)$$

The novel object recognition (NOR) test assessed rat recognition memory and recall of the memory process [4] by measuring the innate exploratory behavior of a novel object in comparison to a familiar one [5]. The test was conducted in a square arena (40 cm × 40 cm) following a three-stage protocol: 1) habituation to the polycarbonate box with wood chips bedding for 10 minutes (40 cm square arena), 2) familiarization: for 3 minutes it was allowed to the rats to get familiarized with two identical glass jars (transparent glass and metal lid) placed in the center of the arena, separated by ~10 cm, and 3) recognition test: one familiar object was replaced with a novel object having distinct physical characteristics (transparent glass light bulb with a metal cap). Then, rats were allowed to explore the arena for 3 minutes. The locations of familiar and novel objects were counterbalanced across trials (left/right). All objects were cleaned with 70% ethanol between each phase. Individual rats were video recorded with an HD camera (model S4612) during the familiarization and recognition test phases.

Videos were preprocessed using ffmpeg to ensure consistent dimensions and object positions. Similar to the EPM test, automated behavioral tracking was performed using DeepLabCut v. 2.3.8. Two separate neural networks were trained, one for each test phase (familiarization and recognition). Each network was trained using 10 distinct frames selected from 10 videos of different animals per phase. New videos were continually added, necessitating re-labeling and re-training. As with the EPM analysis, a ResNet-50 based neural network with default parameters was used for training until a total of 1,620,000 iterations was reached. The recognition trained network got a test error of 3 pixels and a train error of 3 pixels. With a p-cutoff of 0.6, a train error of 2.74 pixels, and a test error of 3.06 pixels. Data tracking generated by DeepLabCut was further processed using a custom DLCAnalyzer script. Time spent exploring familiar and novel objects was used to calculate the discrimination ratio (Eq. 3).

Eq. 3:

$$Discrimination\ ratio\ = \frac{Recognition\ time\ to\ novel\ object\ (seconds)}{Recognition\ time\ to\ novel\ object\ (s) + Recognition\ time\ to\ familiar\ object\ (s)}$$

The conditioned place preference (CPP) is a preclinical modeling paradigm aimed at studying the rewarding and aversive effects of substances like ethanol [6,7]. The apparatus consisted of a three-chambered box with distinct visual and tactile features (white walls/rough floor with ethanol access, gray walls/smooth floor, black walls/smooth floor with water access). The test evaluated the animal's compartment preference across three stages: 1) pre-test (baseline time spent in each compartment without any substance), 2) habituation (6 days to eliminate novelty and for chamber-stimulus association), and 3) test (evaluation of compartment time spent). The preference index was calculated using the time (in seconds) in each chamber (Eq. 4).

Eq. 4:

$$Preference\ index\ = \frac{EtOH\ chamber\ time}{Total\ time\ on\ EtOH\ and\ H2O\ time}$$

Labels used to train the deeplabcut networks.

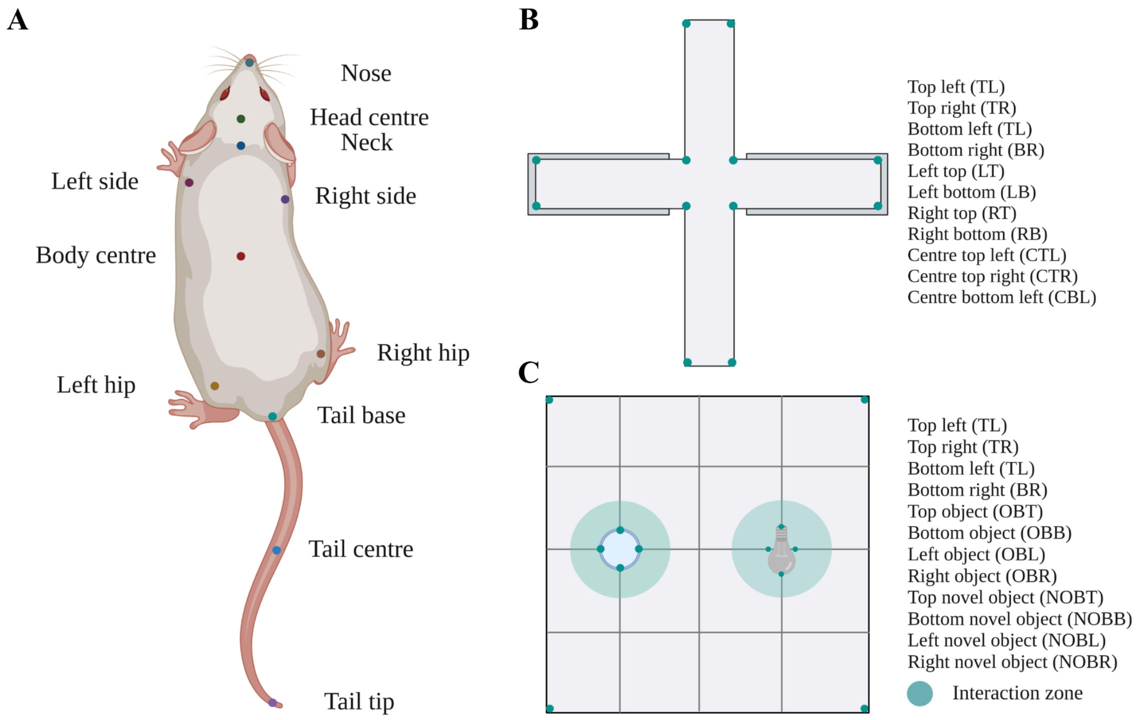


A) The points of interest used to track the rat in each maze, B) The points of interest in the novel object recognition test, C) The points of interest in the elevated plus maze.

### Magnetic Resonance Imaging (MRI) acquisition

Neuroimaging data was obtained at the National Laboratory for Magnetic Resonance Imaging (LANIREM) using a 7 Tesla Magnetic Resonance Imaging (MRI) scanner (Bruker Pharmascan 70/16 US) with a 2 x 2 array surface rat head coil. Structural MRI (sMRI) was acquired with a 3D FLASH sequence T2w with 2 repetitions, TR = 30.76 ms, TE = 5 ms, flip angle = 10º, FOV = 25.6 x 19.098 x 25.6 mm, and an isometric voxel of 160 microns. For the resting state functional MRI (fMRI), we acquired a Gradient-echo Echo-planar imaging (GE EPI) sequence with the following parameters TR= 1,000 ms, TE= 20 ms, flip angle = 60º, slice thickness = 1 mm, FOV = 30 x 30, number of slices = 24, volumes = 600. The interface software used was Paravision v.7.0.

### Structural analysis

Briefly, all individual scans were iteratively averaged between sessions and subjects by using an affine and non-linear approach. The unbiased data-driven study-specific template is generated by iteratively registering each image to the dataset consensus average. The study-specific template created was used to calculate the relative log-transformed Jacobian determinants(46,47) as a measure of brain volume changes, using SIGMA v. 1.2.1 as the template atlas.(45) Then we calculated the individual volume deformations at the voxel level by non-linearly registering each individual image to the template space and calculated the relative Jacobian determinants(46,47) for each subject and session.

The partial least squares (PLS) correlation maximizes the associations between the structural or functional MRI data and the behavioral measures by using the singular value decomposition (SVD) algorithm (details of PLS can be found in (46–48) which decompose the matrices into two matrices of orthonormal singular vectors (U = MRI matrix, and V = behavior matrix), comprising pairs of latent variables (LV) that represents the maximum covariance. A permutation test (1,000 iterations) on the singular values was performed to assess the significance of the LVs.

### Functional analysis

Briefly, the preprocessing included motion correction via rigid registration. Structural images enabled both fMRI-to-anatomical and unbiased template registration. An unbiased template (0.3 mm isotropic voxels) in standard space was constructed, followed by nonlinear registration to the SIGMA v. 1.2.1 atlas. EPI susceptibility distortions were addressed with inhomogeneity correction (adjusting the signal intensity to make it more uniform) before registration for the anatomical scan. Additional steps included slice timing correction, temporal smoothing, denoising, ICA-AROMA for artifact removal, white-matter (WM), cerebrospinal fluid (CSF), head motion translation, and rotation parameters as nuisance regressors. Preprocessed images underwent quality control, and one scan (1 subject-1 session) was excluded.

The functional analysis involved extracting connectivity matrices from the brain regions defined by SIGMA parcellation and using NBS with 1000 random permutations as the null distribution of the strength of connectivity (maximum statistic). This permutation approach helps to determine the statistical significance of the observed differences while controlling for chance findings. The mask used was the same as the one used in structural analysis, with shrinking edges to remove the colliding of the time series.

### MRI anesthesia protocol

For the MRI acquisition, all rats were anesthetized outside the scanner, as reported in Lopez-Castro, et al.[41] The unconsciousness was reached by the induction of anesthesia with vaporized isoflurane ~4% in a transparent polypropylene chamber. Rats were placed in a supine position at the trail of the scanner and were injected with dexmedetomidine hydrochloride (Dexdomitor vet, Zoetis Inc) sterile injectable solution. The solution was administered subcutaneously at a dose of 0.012 mg/kg in a 1 mL bolus under the skin of the interscapular area tented by the thumb and index finger. Time was taken at the moment of dexmedetomidine injection to ensure that the fMRI acquisition was performed after ~30 minutes. Anesthesia was maintained once the rats were in the trail scanner. The composition was a 50/50 mixture of isoflurane/oxygen administered through airflow to the nose, which ranged from 0.6 to 1% of isoflurane. Cardiac and respiratory rates were monitored at all times to prevent distress and spontaneous wake-ups during the MRI acquisitions.

### Statistical analysis

We evaluated the effect of chronic stress and/or ethanol intake with relative volume, functional correlation, weight change, ethanol intake, anxiety index, and preference index as dependent variables in different models (Eq. 5). Group, session, and sex were included as interaction fixed effects (independent variables). In contrast, the discrimination ratio was analyzed using a linear regression model for the NOR task (Eq. 6). Batch was used as a covariate in all models and rat identification (RID) as a random effect. Standardized parameters were obtained by fitting the model on a standardized version of the dataset, 95% confidence intervals (CIs) and p-values were computed using a Wald t-distribution approximation.

To assess group differences in MRI analyses, linear mixed-effects models were applied (Eq. 7). Batch served as a covariate in all models, while group, age, and sex were included as interaction fixed effects.

Eq. 5:

$$Metric \sim Group{\cdot}Age{\cdot}Sex\ +\ Group\ +\ Age\ +\ Sex\ +\ Batch\ \ +\ (1|RID)$$

Metrics: local volume, functional connectivity, ethanol main intake, Δweight = (weight in each session - weight of the first session), anxiety index, and preference index.

Eq. 6:

$$Metric \sim Group{\cdot}Sex\ +\ Group\ +\ Sex\ +\ Batch$$

Metrics: discrimination ratio.

7. Prus AJ, James JR, Rosecrans JA. Conditioned Place Preference. In: Buccafusco JJ, ed. *Methods of Behavior Analysis in Neuroscience*. CRC Press/Taylor & Francis. https://www.ncbi.nlm.nih.gov/pubmed/21204336
