## Supplementary results for "The effect of chronic stress and chronic alcohol intake on behavior, brain structure, and functional connectivity in a rat model"

#### Behavioral test

Individual EPM metric analysis showed that ethanol female groups (EtOH+/CRS- and EtOH+/CRS+), compared to the female EtOH-/CRS- group, spent less percentage time moving ($p_{fdr} = 0.006$, $p_{fdr} = 0.004$) with less speed ($p_{fdr} = 0.031$, $p_{fdr} = 0.022$), but more stationary time ($p_{fdr} = 0.02$, $p_{fdr} = 0.013$) and total spent time ($p_{fdr} = 0.04$, $p_{fdr} = 0.029$) in the closed top arm. CRS male groups (EtOH-/CRS+ and EtOH+/CRS+) only showed significant results compared with the male EtOH-/CRS- group. The EtOH-/CRS+ males spent more time moving ($p_{fdr} = 0.037$), specifically in the center ($p_{fdr} = 0.015$), and spent more time ($p_{fdr} = 0.044$) in the closed bottom arm. Compared with male control group, the male EtOH+/CRS+ group moved during more time ($p_{fdr} = 0.012$) with more distance covered ($p_{fdr} = 0.009$), more transitions into ($p_{fdr} = 0.04$), and spending more time ($p_{fdr} = 0.048$) in the center of the arm, but more stationary time ($p_{fdr} = 0.048$) in both closed arms. Less transitions ($p_{fdr} = 0.027$) to the closed top arm, more distance covered ($p_{fdr} = 0.034$), more transitions into ($p_{fdr} = 0.014$), time ($p_{fdr} = 0.03$) and distance moving ($p_{fdr} = 0.03$) in the left open arm, and less transitions ($p_{fdr} = 0.025$) in the open right. This group also showed more transitions to the left ($p_{fdr} = 0.025$, $p_{fdr} = 0.002$) and right arm ($p_{fdr} = 0.009$, $p_{fdr} = 0.029$).

Discrimination ratio of the NOR task (Eq. 3, supplementary results) showed a lower index only in the male EtOH-/CRS+ group compared with male EtOH+/CRS- ($\beta = 0.26$, $p_{fdr} = 0.023$, $p = 0.003$, $d = 1.53$), and uncorrected significant compared with male EtOH-/CRS- ($\beta = 0.19$, $p_{fdr} = 0.096$, $p = 0.048$, $d = 1.1$) and with male EtOH+/CRS+ ($\beta = -0.17$, $p_{fdr} = 0.096$, $p = 0.048$, $d = -0.97$) (Figure 3d). In particular, this group spent less time in the novel object ($p_{fdr} = 0.044$) with less stationary time ($p_{fdr} = 0.037$) and speed moving ($p_{fdr} = 0.046$). Interestingly, the male EtOH+/CRS+ group moved more distance ($p_{fdr} = 0.026$) with higher speed ($p_{fdr} < 0.001$) compared with male EtOH+/CRS-, into the familiar object (speed moving: $p_{fdr} = 0.012$), in the novel one (speed moving: $p_{fdr} = 0.028$), and showed higher speed compared with EtOH-/CRS- in familiar ($p_{fdr} = 0.019$) and novel ($p_{fdr} = 0.039$) objects. In contrast, females EtOH-/CRS+, compared with EtOH-/CRS- of the same sex, showed higher time spent ($p_{fdr} = 0.025$), distance moved ($p_{fdr} = 0.04$) and moving time ($p_{fdr} = 0.011$) in the familiar object. In the similar way that males, the EtOH+/CRS+ female group showed more distance moving ($p_{fdr} = 0.025$) compared with EtOH+/CRS-, into the familiar object (speed moving: $p_{fdr} = 0.04$) and in the novel one (speed moving: $p_{fdr} = 0.015$). Besides, this contrast also showed spending more stationary time ($p_{fdr} = 0.04$) and transitions in both object (familiar: $p_{fdr} = 0.035$, novel: $p_{fdr} = 0.025$) and distance moving (familiar $p_{fdr} = 0.015$, novel: $p_{fdr} = 0.015$). Besides, female EtOH+/CRS+ contrasted to EtOH-/CRS- group had more distance moving ($p_{fdr} = 0.004$), total time spent ($p_{fdr} = 0.022$) and transitions ($p_{fdr} = 0.001$) in familiar object, but more distance moving ($p_{fdr} = 0.015$) and time moving ($p_{fdr} = 0.048$) in only novel object.

#### Voxelwise analysis of each group contrasted with the control

Ethanol intake groups (EtOH+/CRS- and EtOH+/CRS+) exhibited an enlargement in the entorhinal cortex (Ent) and thalamus (Thal), along with a reduction in the hypothalamus (Hyp) and cerebellum. Conversely, chronic restraint groups (EtOH-/CRS+ and EtOH+/CRS+) showed an increase solely in the insular cortex (Ins) and a decrease in the thalamus, secondary motor cortex (M2), and retrosplenial cortex (RSG). The EtOH+/CRS- group alone showed volume alterations in the substantia nigra (decrease) and perirhinal cortex (increase). The EtOH-/CRS+ group displayed an increase in the orbitofrontal lobule (Orb), while EtOH-/CRS+ exhibited an expansion in the secondary cingulate cortex (Cg2) and a reduction in the secondary motor cortex (M2). The amygdala (Amy) region showed opposite volume changes between EtOH+/CRS- (higher) and EtOH-/CRS+ (lower) compared to the control group.

Compared to the only stress (EtOH-/CRS+) group, ethanol intake groups exhibited increased volumes in the olfactory bulb (OB), cerebellum (Cer), entorhinal cortex (MEntR), and thalamus (Thal), alongside decreased volumes in the dorsal caudate-putamen (CPu) and Thal. Additional regions, such as the nucleus accumbens (Nacc), hippocampus (Hipp), insular cortex (Ins), and hypothalamus (Hyp), showed increased volumes in the EtOH+/CRS- group but decreased volumes in the EtOH+/CRS+ group. The substantia nigra (SN) and secondary motor cortex (M2) displayed increased volumes, whereas the frontal cortex, secondary cingulate cortex (Cg2), and amygdala (Amy) exhibited decreased volumes. The orbitofrontal cortex (Orb), retrosplenial cortex (RSG), and primary motor cortex (M1) showed reduced volumes exclusively in the EtOH+/CRS+ group. In contrast to ethanol intake (EtOH+/CRS-), chronic restraint stress led to increased volumes in the OB, Cer, Cg2, MEntR, CPu, and Thal in both groups (EtOH+/CRS+ and EtOH-/CRS+). This increase was accompanied by a decrease in the volumes of the Hipp (DG), Thal, and Amy.

### Supplementary tables

**Supplementary table 1.** Group pairwise comparison of longitudinal ROI functional connectivity changes based on estimated marginal means.

| Contrast | | ROI | ROI | Sex | Mean | d | SE | df | p-value | q-value |
|---|---|---|---|---|---|---|---|---|---|---|
| EtOH-/CRS- | EtOH+/CRS- | rAmy | lvCer | fem | 0.08 | 0.82 | 0.02 | 77 | < 0.001 | 0.01 |
| | | rCg2 | rThal | fem | -0.09 | -0.83 | 0.03 | 77 | < 0.001 | < 0.001 |
| | EtOH-/CRS+ | rAmy | lThal | fem | 0.1 | 0.86 | 0.04 | 80 | 0.01 | 0.04 |
| | | rCg2 | rThal | fem | -0.12 | -1.04 | 0.03 | 80 | < 0.001 | < 0.001 |
| | EtOH+/CRS+ | rCg2 | lM2 | fem | -0.11 | -0.64 | 0.04 | 82 | 0.01 | 0.049 |
| | | rCg2 | rThal | fem | -0.15 | -1.04 | 0.03 | 81 | < 0.001 | < 0.001 |
| EtOH+/CRS- | EtOH-/CRS+ | rHipp | rvCer | fem | 0.08 | 0.77 | 0.03 | 78 | 0.01 | 0.04 |
| | EtOH+/CRS+ | lCg1 | lStr | male | 0.1 | -0.21 | 0.03 | 81 | 0.01 | 0.03 |
| EtOH-/CRS+ | EtOH+/CRS+ | rHipp | rvCer | fem | -0.09 | 0.69 | 0.03 | 74 | < 0.001 | 0.02 |
| | | rEnt | lSN | male | -0.08 | -0.56 | 0.03 | 76 | < 0.001 | 0.03 |

Abbreviation: Mean = estimated marginal means, d = effect size, SE = standard error, df = degree of freedom, Q-value = p-value fdr adjusted.

### Supplementary figures

**Supplementary figure 1.** Regional volume trajectory changes of each group contrasted to EtOH-/CRS-

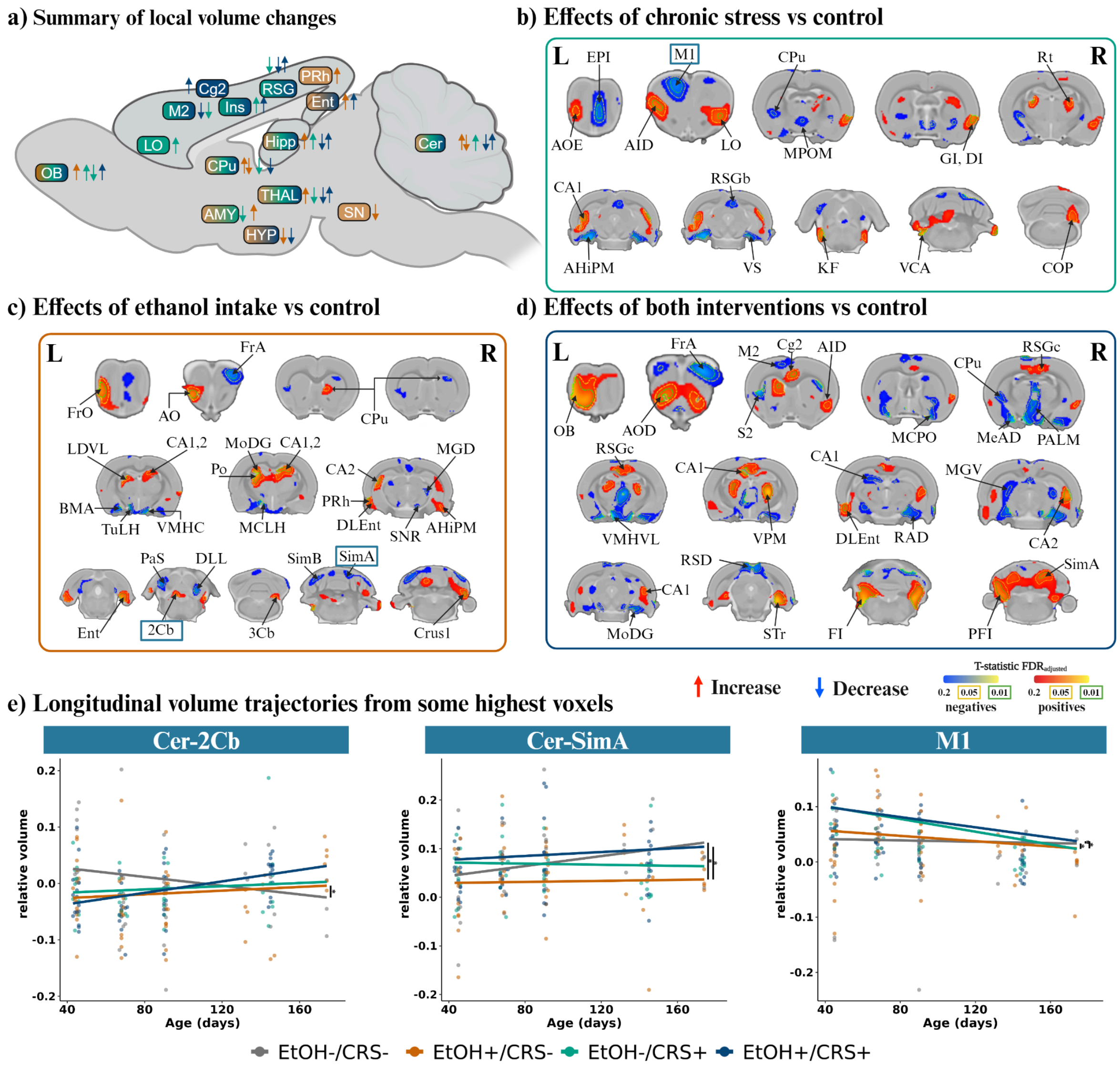


**a)** Schematic summary of the regional volume changes found on each contrast. **b)** Volume differences associated with stress effects (EtOH-/CRS+). **c)** Volume differences associated with ethanol effects (EtOH+/CRS-). **d)** Volume differences associated with stress and ethanol effects (EtOH+/CRS+). **e)** Scatter plot of the corrected volume with regression lines of the voxel with highest t-value of the ROI highlighted with a box (See table 3). Threshold set at FDR of 20%, 5% (yellow dashed line) and 1% (green dashed line). Coronal slides labels correspond to Paxinos & Watson stereotaxic atlas.

**Supplementary figure 2.** regional volume trajectory changes of each group contrasted to EtOH-/CRS+

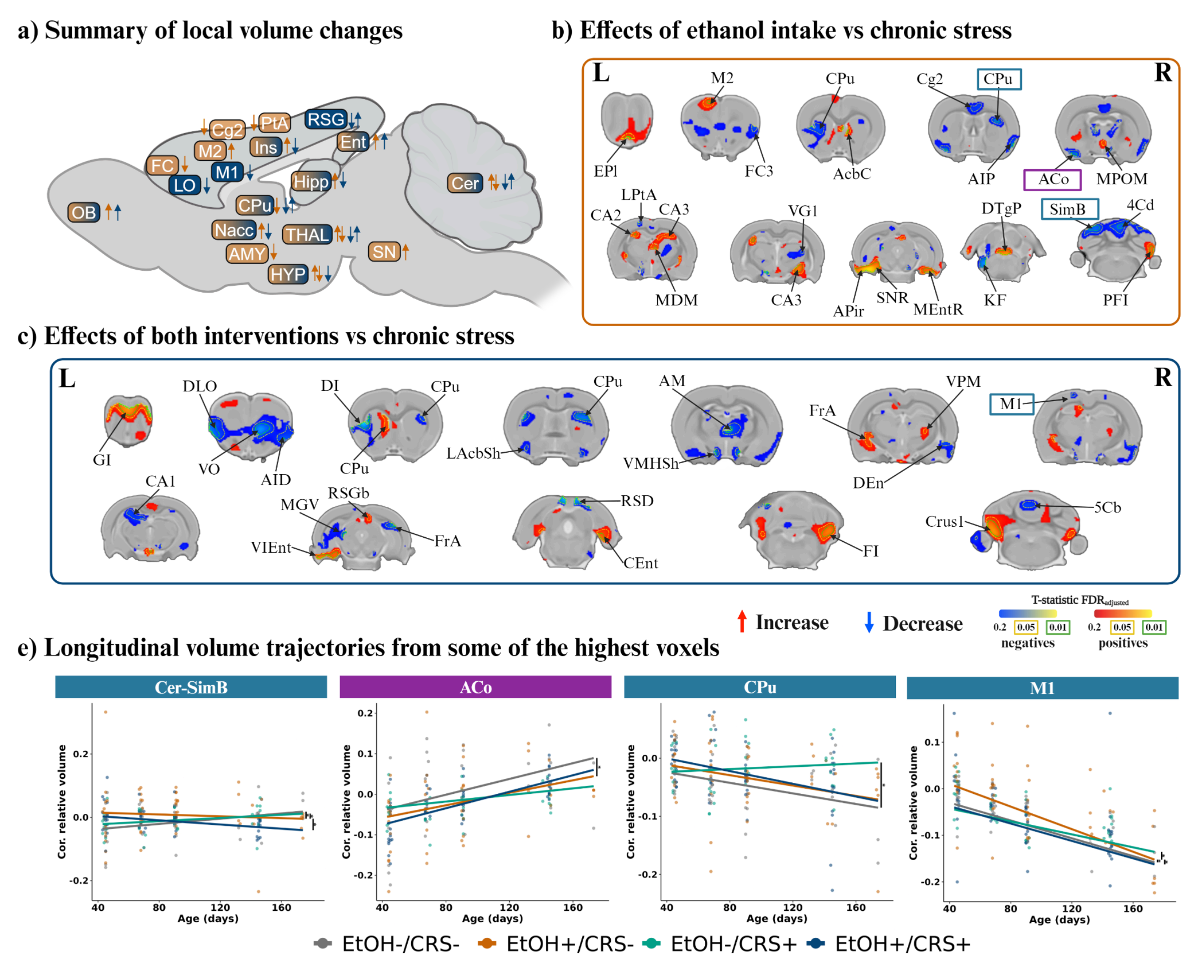


**a)** Schematic summary of the regional volume changes found on each contrast. **b)** Volume differences associated with ethanol effects without stress (EtOH+/CRS-). **c)** Volume differences associated with just ethanol effects (EtOH+/CRS+). **e)** Scatter plot of the corrected volume with regression lines of the voxel with highest t-value of the ROI highlighted with a box (See table 3). Threshold set at FDR of 20%, 5% (yellow dashed line) and 1% (green dashed line). Coronal slides labels correspond to Paxinos & Watson stereotaxic atlas.

**Supplementary figure 3.** Regional volume trajectory changes of each group compared to EtOH+/CRS-

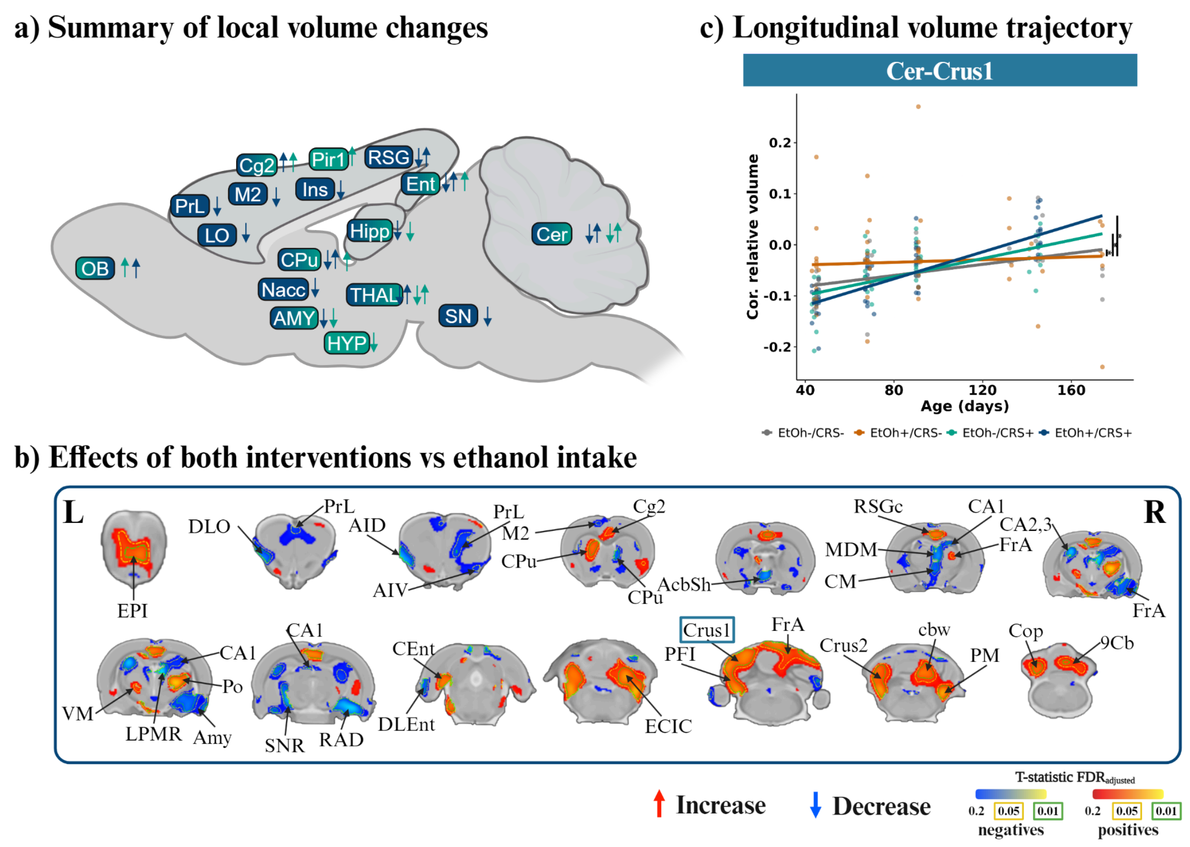


**a)** Schematic summary of the regional volume changes found on each contrast. **b)** Volume differences associated with just stress effects (EtOH+/CRS+). **c)** Scatter plot of the corrected volume with regression lines of the voxel with highest t-value (See table 3). Threshold set at FDR of 20%, 5% (yellow dashed line) and 1% (green dashed line). Coronal slides labels correspond to Paxinos & Watson stereotaxic atlas.

**Supplementary figure 4.** Corticosterone levels from a sample of each group at the end of the protocol (P142).

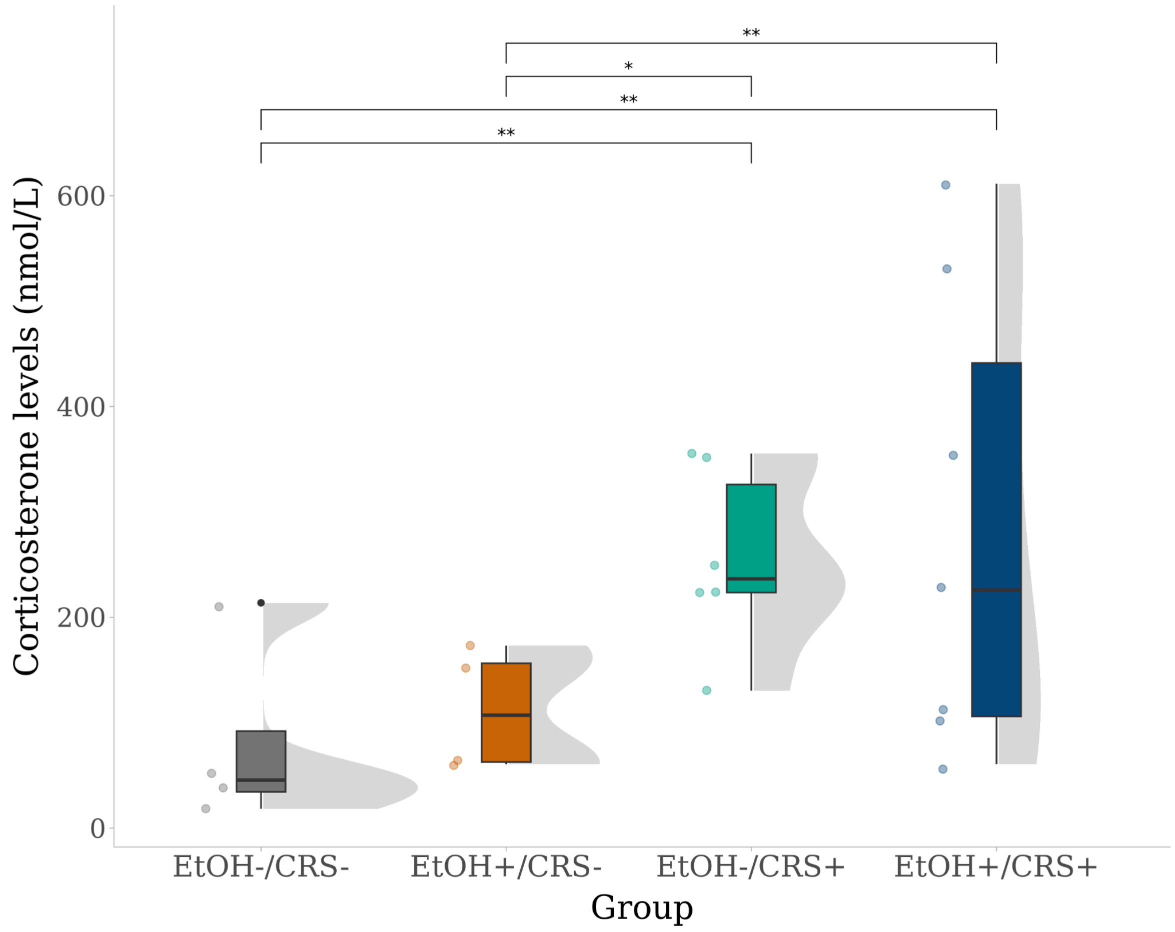


Corticosterone levels of EtOH-/CRS- (n = 4), EtOH+/CRS- (n = 4), EtOH-/CRS+ (n = 6), EtOH+/CRS+ (n = 7).

### Individual metrics of elevated plus maze (EPM) and their group differences

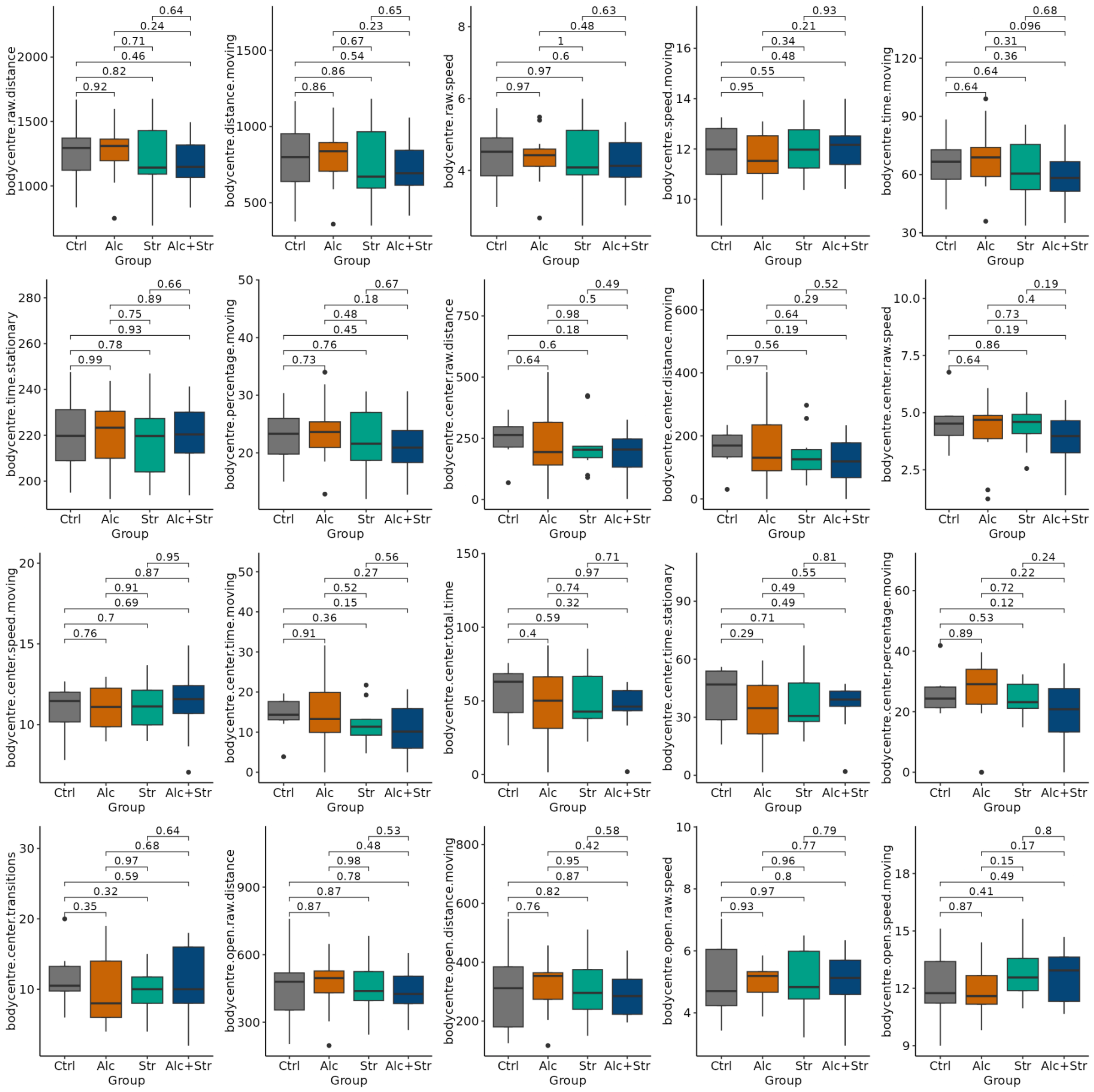

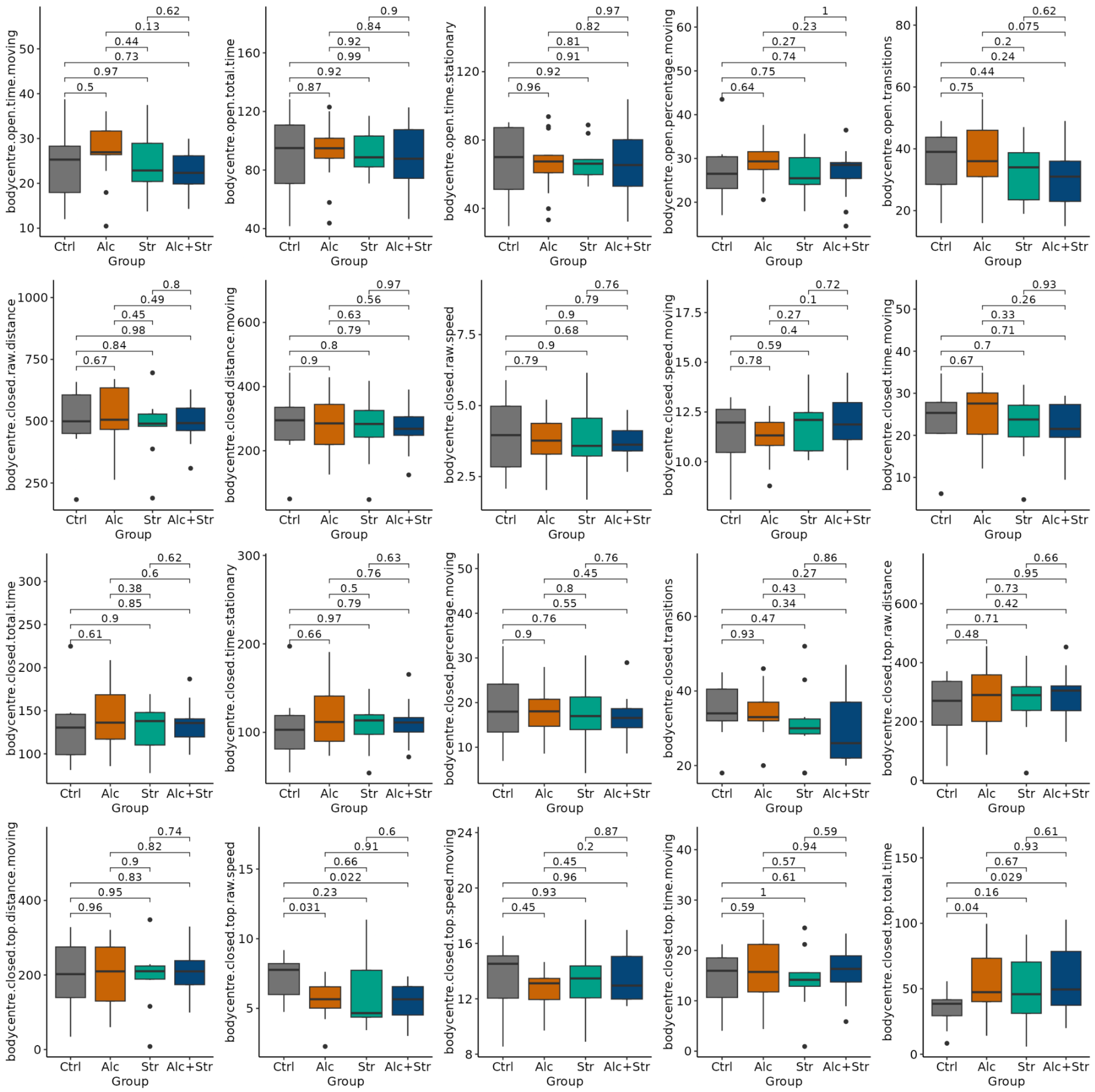

bodycentre.open.time.moving
0.62 0.13 0.44 0.73 0.97 0.5
bodycentre.open.total.time
0.9 0.84 0.92 0.99 0.92 0.87
bodycentre.open.time.stationary
0.97 0.82 0.81 0.91 0.92 0.96
bodycentre.open.percentage.moving
1 0.23 0.27 0.74 0.75 0.64
bodycentre.open.transitions
0.62 0.075 0.2 0.24 0.44 0.75
bodycentre.closed.raw.distance
0.8 0.49 0.45 0.98 0.84 0.67
bodycentre.closed.distance.moving
0.97 0.56 0.63 0.79 0.8 0.9
bodycentre.closed.raw.speed
0.76 0.79 0.9 0.68 0.9 0.79
bodycentre.closed.speed.moving
0.72 0.1 0.27 0.4 0.59 0.78
bodycentre.closed.time.moving
0.93 0.26 0.33 0.71 0.7 0.67
bodycentre.closed.total.time
0.62 0.6 0.38 0.85 0.9 0.61
bodycentre.closed.time.stationary
0.63 0.76 0.5 0.79 0.97 0.66
bodycentre.closed.percentage.moving
0.76 0.45 0.8 0.55 0.76 0.9
bodycentre.closed.transitions
0.86 0.27 0.43 0.34 0.47 0.93
bodycentre.closed.top.raw.distance
0.66 0.95 0.73 0.42 0.71 0.48
bodycentre.closed.top.distance.moving
0.74 0.82 0.9 0.83 0.95 0.96
bodycentre.closed.top.raw.speed
0.6 0.91 0.66 0.022 0.23 0.031
bodycentre.closed.top.speed.moving
0.87 0.2 0.45 0.96 0.93 0.45
bodycentre.closed.top.time.moving
0.59 0.94 0.57 0.61 1 0.59
bodycentre.closed.top.total.time
0.61 0.93 0.67 0.029 0.16 0.04
Ctrl Alc Str Alc+Str
Group

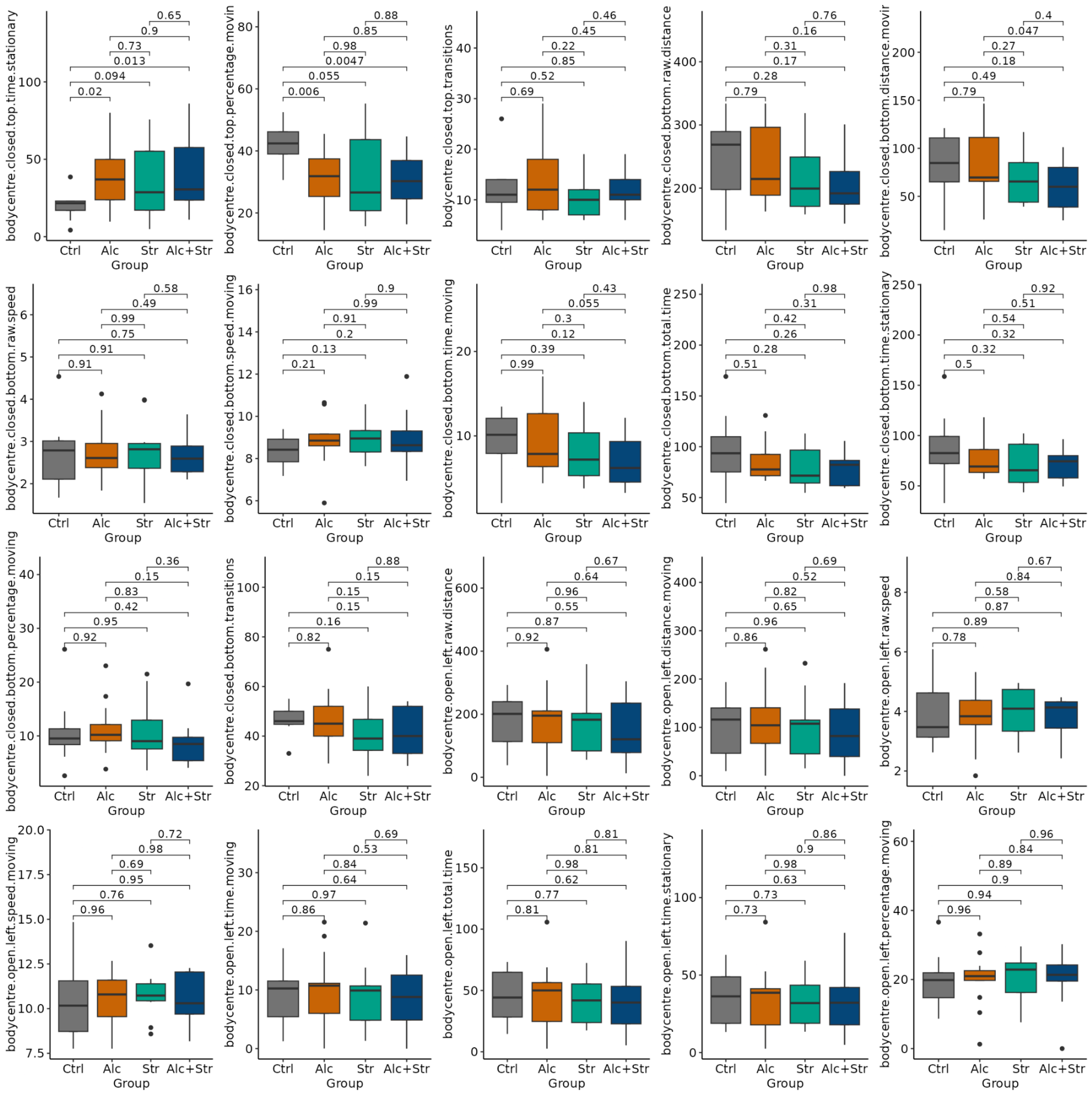
bodycentre.closed.top.time.stationary
0.65 0.9 0.73 0.013 0.094 0.02
bodycentre.closed.top.percentage.movin
0.88 0.85 0.98 0.0047 0.055 0.006
bodycentre.closed.top.transitions
0.46 0.45 0.22 0.85 0.52 0.69
bodycentre.closed.bottom.raw.distance
0.76 0.16 0.31 0.17 0.28 0.79
bodycentre.closed.bottom.distance.movir
0.4 0.047 0.27 0.18 0.49 0.79
bodycentre.closed.bottom.raw.speed
0.58 0.49 0.99 0.75 0.91 0.91
bodycentre.closed.bottom.speed.moving
0.9 0.99 0.91 0.2 0.13 0.21
bodycentre.closed.bottom.time.moving
0.43 0.055 0.3 0.12 0.39 0.99
bodycentre.closed.bottom.total.time
0.98 0.31 0.42 0.26 0.28 0.51
bodycentre.closed.bottom.time.stationary
0.92 0.51 0.54 0.32 0.32 0.5
bodycentre.closed.bottom.percentage.moving
0.36 0.15 0.83 0.42 0.95 0.92
bodycentre.closed.bottom.transitions
0.88 0.15 0.15 0.15 0.16 0.82
bodycentre.open.left.raw.distance
0.67 0.64 0.96 0.55 0.87 0.92
bodycentre.open.left.distance.moving
0.69 0.52 0.82 0.65 0.96 0.86
bodycentre.open.left.raw.speed
0.67 0.84 0.58 0.87 0.89 0.78
bodycentre.open.left.speed.moving
0.72 0.98 0.69 0.95 0.76 0.96
bodycentre.open.left.time.moving
0.69 0.53 0.84 0.64 0.97 0.86
bodycentre.open.left.total.time
0.81 0.81 0.98 0.62 0.77 0.81
bodycentre.open.left.time.stationary
0.86 0.9 0.98 0.63 0.73 0.73
bodycentre.open.left.percentage.moving
0.96 0.84 0.89 0.9 0.94 0.96
Ctrl Alc Str Alc+Str
Group

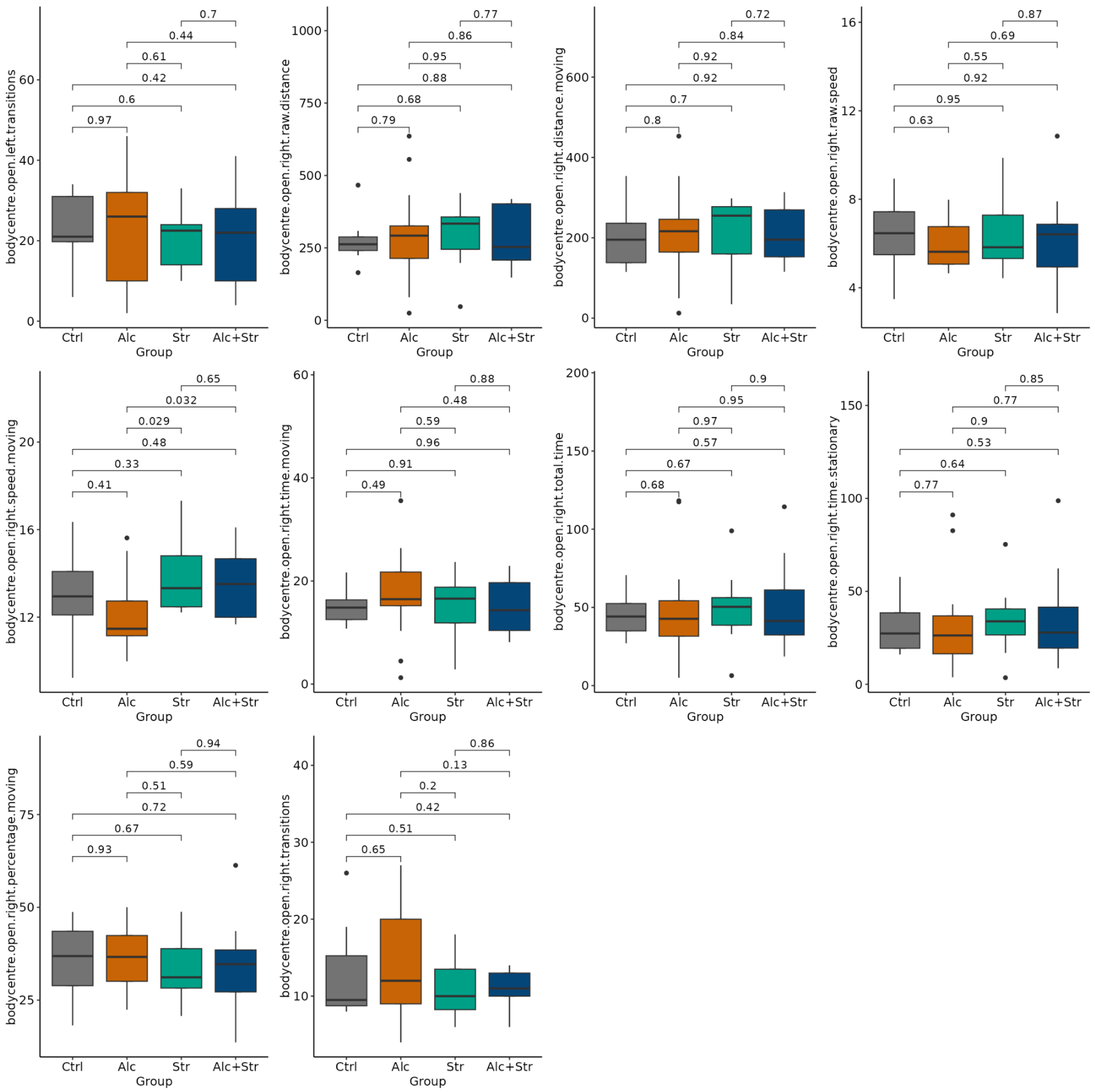
bodycentre.open.left.transitions
0.7
0.44
0.61
0.42
0.6
0.97
0
20
40
60
Ctrl
Alc
Str
Alc+Str
Group
bodycentre.open.right.raw.distance
0.77
0.86
0.95
0.88
0.68
0.79
0
250
500
750
1000
Ctrl
Alc
Str
Alc+Str
Group
bodycentre.open.right.distance.moving
0.72
0.84
0.92
0.92
0.7
0.8
0
200
400
600
Ctrl
Alc
Str
Alc+Str
Group
bodycentre.open.right.raw.speed
0.87
0.69
0.55
0.92
0.95
0.63
4
8
12
16
Ctrl
Alc
Str
Alc+Str
Group
bodycentre.open.right.speed.moving
0.65
0.032
0.029
0.48
0.33
0.41
12
16
20
Ctrl
Alc
Str
Alc+Str
Group
bodycentre.open.right.time.moving
0.88
0.48
0.59
0.96
0.91
0.49
0
20
40
60
Ctrl
Alc
Str
Alc+Str
Group
bodycentre.open.right.total.time
0.9
0.95
0.97
0.57
0.67
0.68
0
50
100
150
200
Ctrl
Alc
Str
Alc+Str
Group
bodycentre.open.right.time.stationary
0.85
0.77
0.9
0.53
0.64
0.77
0
50
100
150
Ctrl
Alc
Str
Alc+Str
Group
bodycentre.open.right.percentage.moving
0.94
0.59
0.51
0.72
0.67
0.93
25
50
75
Ctrl
Alc
Str
Alc+Str
Group
bodycentre.open.right.transitions
0.86
0.13
0.2
0.42
0.51
0.65
10
20
30
40
Ctrl
Alc
Str
Alc+Str
Group

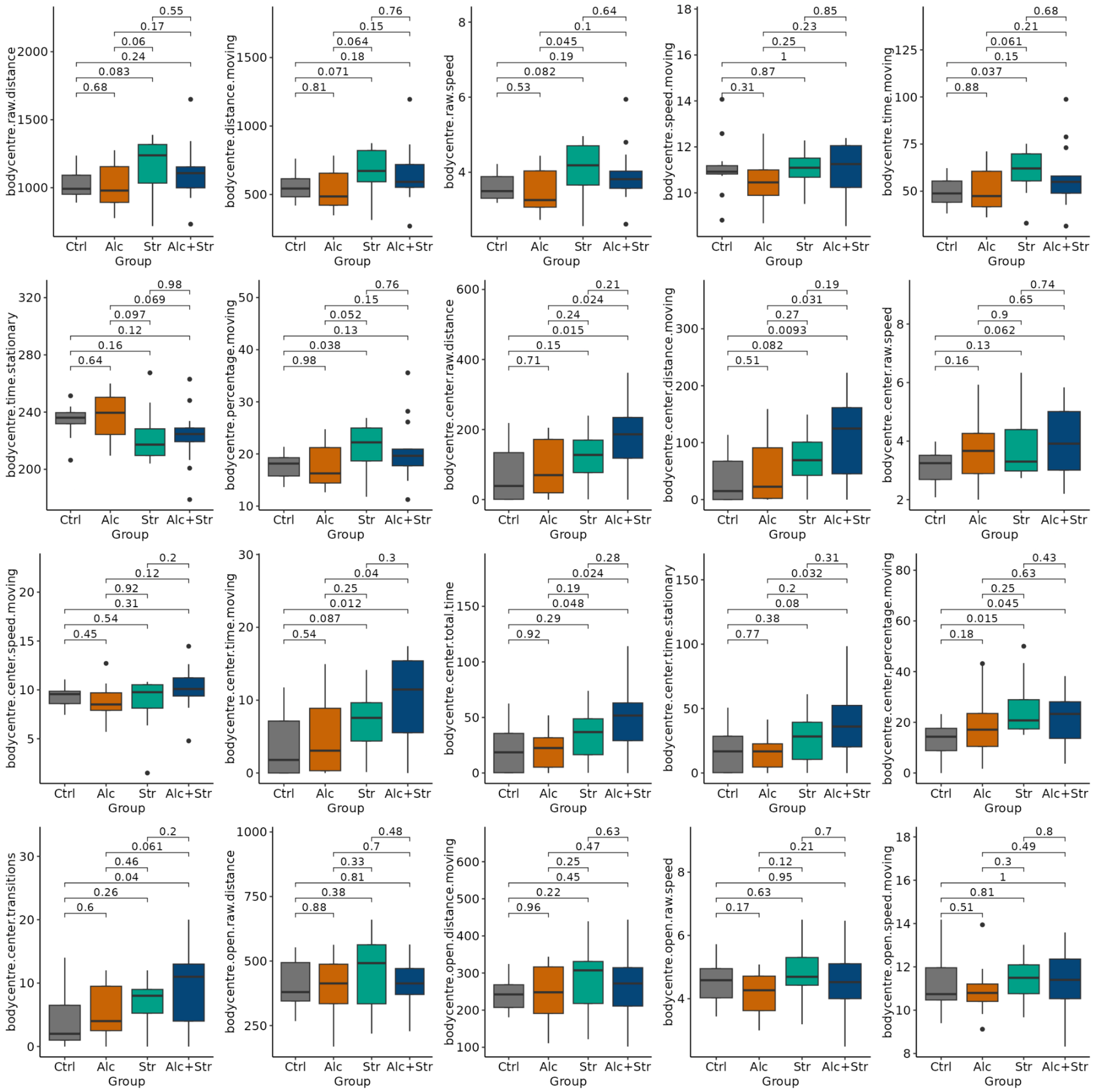

bodycentre.raw.distance
0.55
0.17
0.06
0.24
0.083
0.68
Ctrl
Alc
Str
Alc+Str
Group
bodycentre.distance.moving
0.76
0.15
0.064
0.18
0.071
0.81
bodycentre.raw.speed
0.64
0.1
0.045
0.19
0.082
0.53
bodycentre.speed.moving
0.85
0.23
0.25
1
0.87
0.31
bodycentre.time.moving
0.68
0.21
0.061
0.15
0.037
0.88
bodycentre.time.stationary
0.98
0.069
0.097
0.12
0.16
0.64
bodycentre.percentage.moving
0.76
0.15
0.052
0.13
0.038
0.98
bodycentre.center.raw.distance
0.21
0.024
0.24
0.015
0.15
0.71
bodycentre.center.distance.moving
0.19
0.031
0.27
0.0093
0.082
0.51
bodycentre.center.raw.speed
0.74
0.65
0.9
0.062
0.13
0.16
bodycentre.center.speed.moving
0.2
0.12
0.92
0.31
0.54
0.45
bodycentre.center.time.moving
0.3
0.04
0.25
0.012
0.087
0.54
bodycentre.center.total.time
0.28
0.024
0.19
0.048
0.29
0.92
bodycentre.center.time.stationary
0.31
0.032
0.2
0.08
0.38
0.77
bodycentre.center.percentage.moving
0.43
0.63
0.25
0.045
0.015
0.18
bodycentre.center.transitions
0.2
0.061
0.46
0.04
0.26
0.6
bodycentre.open.raw.distance
0.48
0.7
0.33
0.81
0.38
0.88
bodycentre.open.distance.moving
0.63
0.47
0.25
0.45
0.22
0.96
bodycentre.open.raw.speed
0.7
0.21
0.12
0.95
0.63
0.17
bodycentre.open.speed.moving
0.8
0.49
0.3
1
0.81
0.51

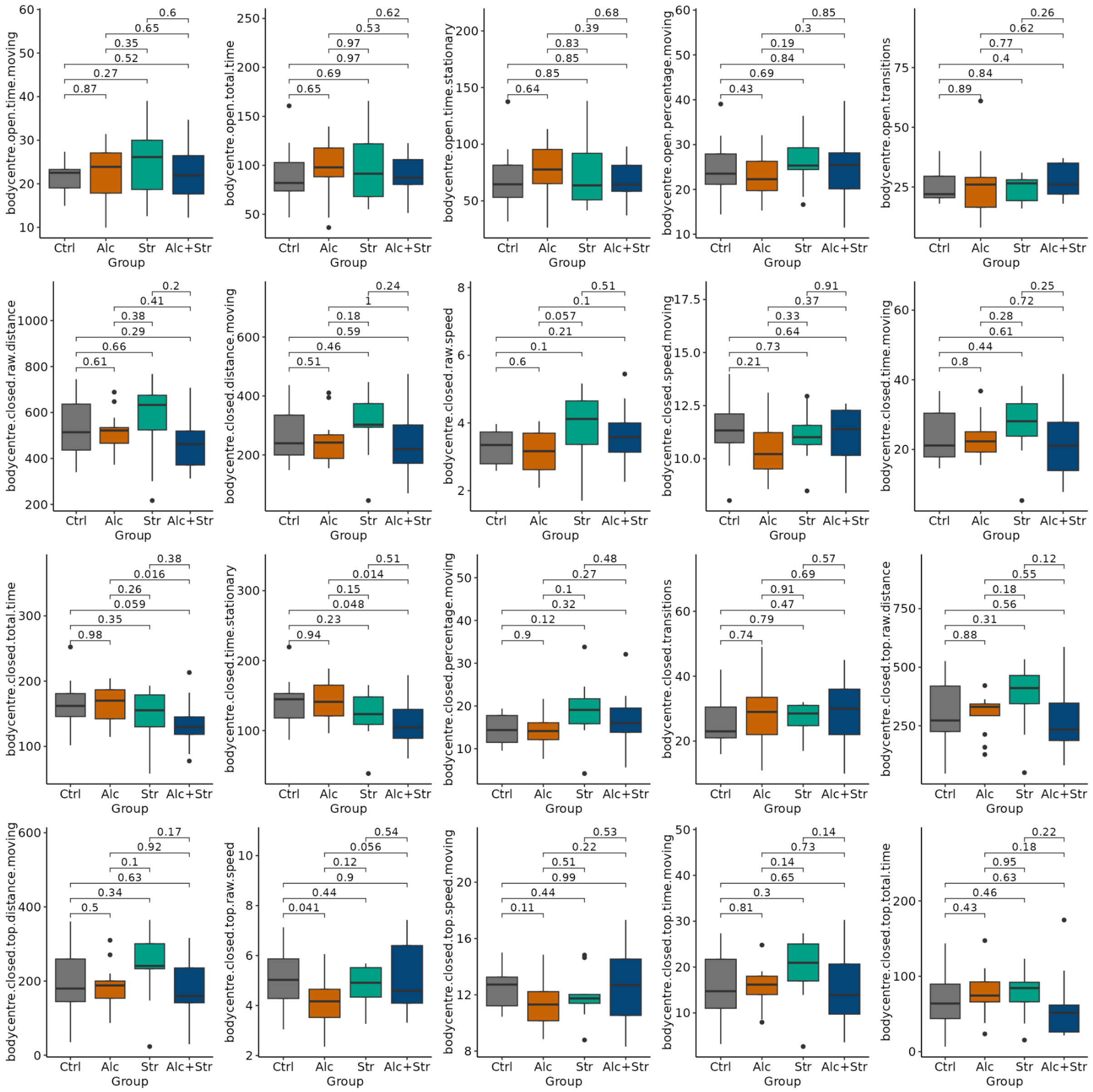

bodycentre.open.time.moving
0.6
0.65
0.35
0.52
0.27
0.87
bodycentre.open.total.time
0.62
0.53
0.97
0.97
0.69
0.65
bodycentre.open.time.stationary
0.68
0.39
0.83
0.85
0.85
0.64
bodycentre.open.percentage.moving
0.85
0.3
0.19
0.84
0.69
0.43
bodycentre.open.transitions
0.26
0.62
0.77
0.4
0.84
0.89
bodycentre.closed.raw.distance
0.2
0.41
0.38
0.29
0.66
0.61
bodycentre.closed.distance.moving
0.24
1
0.18
0.59
0.46
0.51
bodycentre.closed.raw.speed
0.51
0.1
0.057
0.21
0.1
0.6
bodycentre.closed.speed.moving
0.91
0.37
0.33
0.64
0.73
0.21
bodycentre.closed.time.moving
0.25
0.72
0.28
0.61
0.44
0.8
bodycentre.closed.total.time
0.38
0.016
0.26
0.059
0.35
0.98
bodycentre.closed.time.stationary
0.51
0.014
0.15
0.048
0.23
0.94
bodycentre.closed.percentage.moving
0.48
0.27
0.1
0.32
0.12
0.9
bodycentre.closed.transitions
0.57
0.69
0.91
0.47
0.79
0.74
bodycentre.closed.top.raw.distance
0.12
0.55
0.18
0.56
0.31
0.88
bodycentre.closed.top.distance.moving
0.17
0.92
0.1
0.63
0.34
0.5
bodycentre.closed.top.raw.speed
0.54
0.056
0.12
0.9
0.44
0.041
bodycentre.closed.top.speed.moving
0.53
0.22
0.51
0.99
0.44
0.11
bodycentre.closed.top.time.moving
0.14
0.73
0.14
0.65
0.3
0.81
bodycentre.closed.top.total.time
0.22
0.18
0.95
0.63
0.46
0.43
Ctrl
Alc
Str
Alc+Str
Group

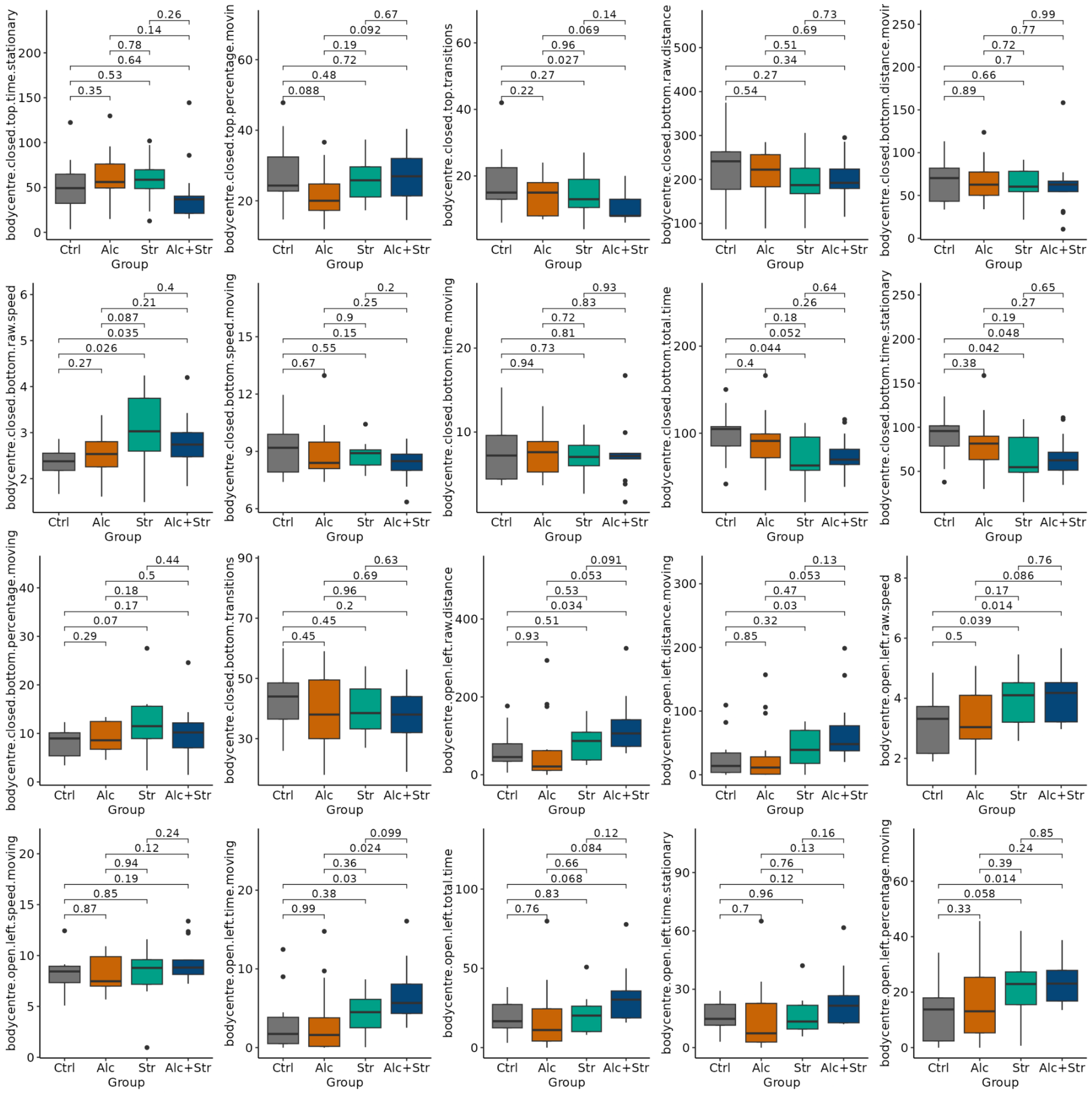
bodycentre.closed.top.time.stationary
0.26
0.14
0.78
0.64
0.53
0.35
bodycentre.closed.top.percentage.movin
0.67
0.092
0.19
0.72
0.48
0.088
bodycentre.closed.top.transitions
0.14
0.069
0.96
0.027
0.27
0.22
bodycentre.closed.bottom.raw.distance
0.73
0.69
0.51
0.34
0.27
0.54
bodycentre.closed.bottom.distance.movir
0.99
0.77
0.72
0.7
0.66
0.89
bodycentre.closed.bottom.raw.speed
0.4
0.21
0.087
0.035
0.026
0.27
bodycentre.closed.bottom.speed.moving
0.2
0.25
0.9
0.15
0.55
0.67
bodycentre.closed.bottom.time.moving
0.93
0.83
0.72
0.81
0.73
0.94
bodycentre.closed.bottom.total.time
0.64
0.26
0.18
0.052
0.044
0.4
bodycentre.closed.bottom.time.stationary
0.65
0.27
0.19
0.048
0.042
0.38
bodycentre.closed.bottom.percentage.moving
0.44
0.5
0.18
0.17
0.07
0.29
bodycentre.closed.bottom.transitions
0.63
0.69
0.96
0.2
0.45
0.45
bodycentre.open.left.raw.distance
0.091
0.053
0.53
0.034
0.51
0.93
bodycentre.open.left.distance.moving
0.13
0.053
0.47
0.03
0.32
0.85
bodycentre.open.left.raw.speed
0.76
0.086
0.17
0.014
0.039
0.5
bodycentre.open.left.speed.moving
0.24
0.12
0.94
0.19
0.85
0.87
bodycentre.open.left.time.moving
0.099
0.024
0.36
0.03
0.38
0.99
bodycentre.open.left.total.time
0.12
0.084
0.66
0.068
0.83
0.76
bodycentre.open.left.time.stationary
0.16
0.13
0.76
0.12
0.96
0.7
bodycentre.open.left.percentage.moving
0.85
0.24
0.39
0.014
0.058
0.33
Ctrl
Alc
Str
Alc+Str
Group

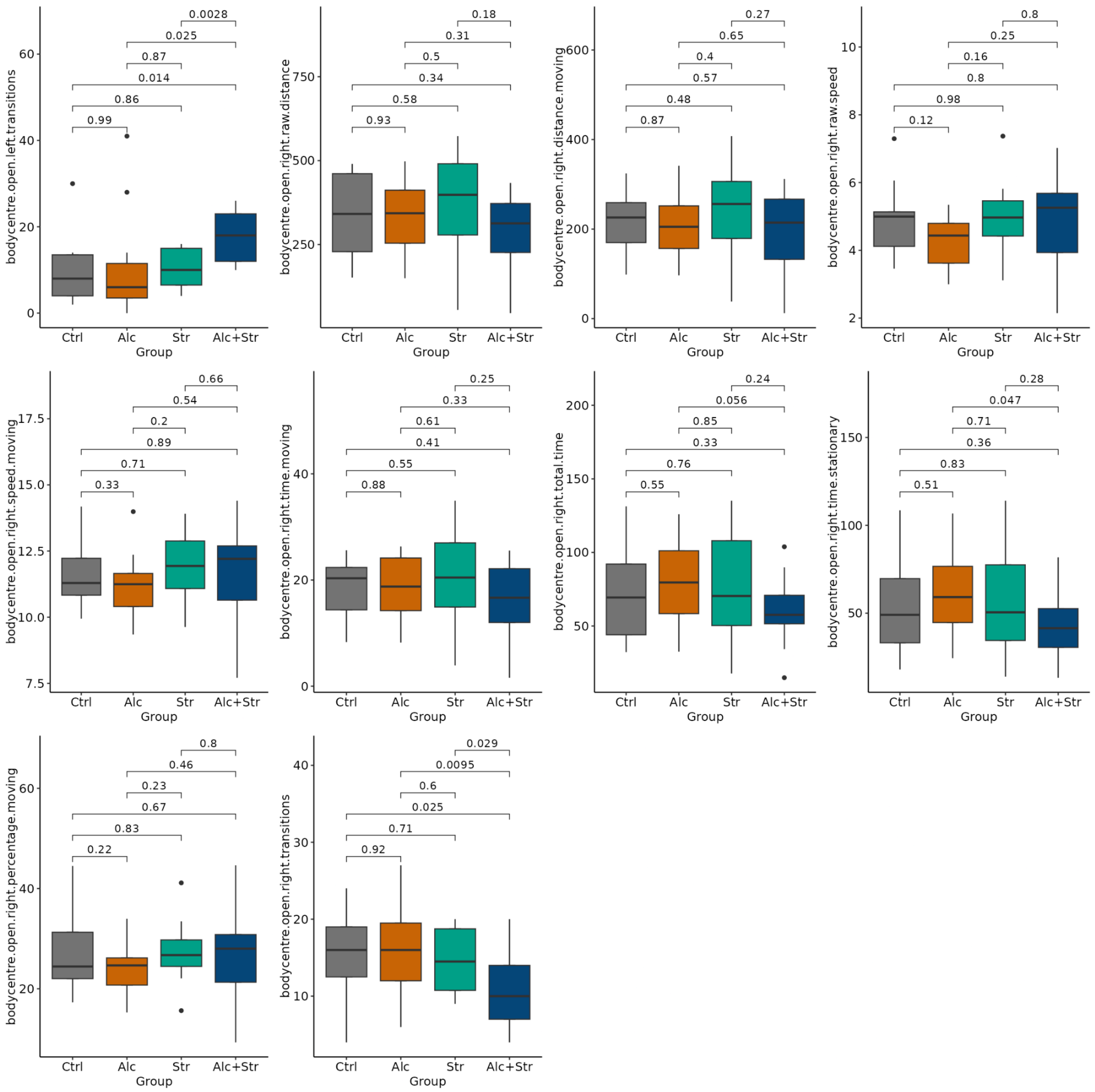
0.0028
0.025
0.87
0.014
0.86
0.99
0
20
40
60
bodycentre.open.left.transitions
Ctrl
Alc
Str
Alc+Str
Group
0.18
0.31
0.5
0.34
0.58
0.93
250
500
750
bodycentre.open.right.raw.distance
Ctrl
Alc
Str
Alc+Str
Group
0.27
0.65
0.4
0.57
0.48
0.87
0
200
400
600
bodycentre.open.right.distance.moving
Ctrl
Alc
Str
Alc+Str
Group
0.8
0.25
0.16
0.8
0.98
0.12
2
4
6
8
10
bodycentre.open.right.raw.speed
Ctrl
Alc
Str
Alc+Str
Group
0.66
0.54
0.2
0.89
0.71
0.33
7.5
10.0
12.5
15.0
17.5
bodycentre.open.right.speed.moving
Ctrl
Alc
Str
Alc+Str
Group
0.25
0.33
0.61
0.41
0.55
0.88
0
20
40
bodycentre.open.right.time.moving
Ctrl
Alc
Str
Alc+Str
Group
0.24
0.056
0.85
0.33
0.76
0.55
50
100
150
200
bodycentre.open.right.total.time
Ctrl
Alc
Str
Alc+Str
Group
0.28
0.047
0.71
0.36
0.83
0.51
50
100
150
bodycentre.open.right.time.stationary
Ctrl
Alc
Str
Alc+Str
Group
0.8
0.46
0.23
0.67
0.83
0.22
20
40
60
bodycentre.open.right.percentage.moving
Ctrl
Alc
Str
Alc+Str
Group
0.029
0.0095
0.6
0.025
0.71
0.92
10
20
30
40
bodycentre.open.right.transitions
Ctrl
Alc
Str
Alc+Str
Group

### Individual metrics of novel object recognition (NOR) and their group differences

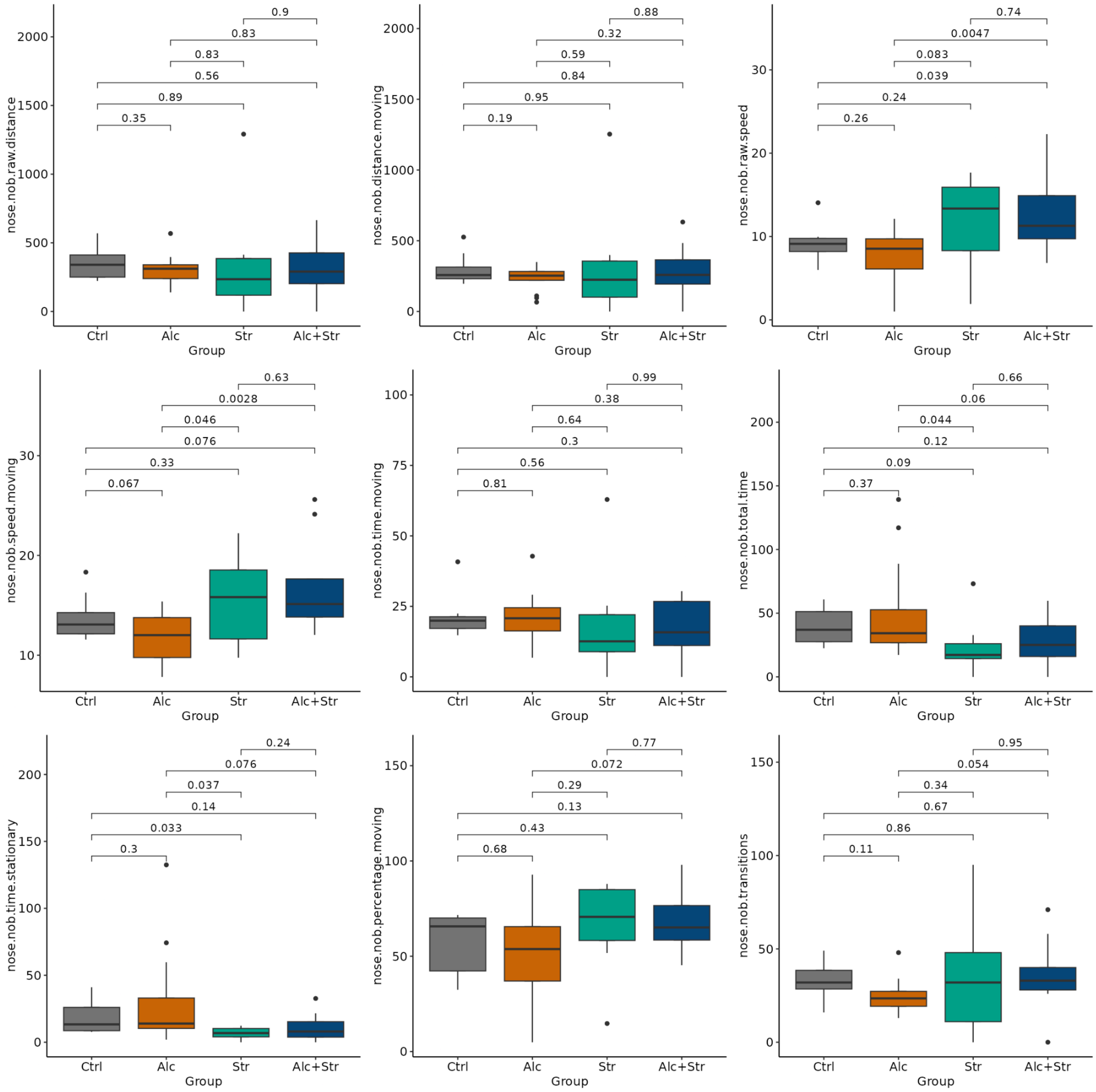

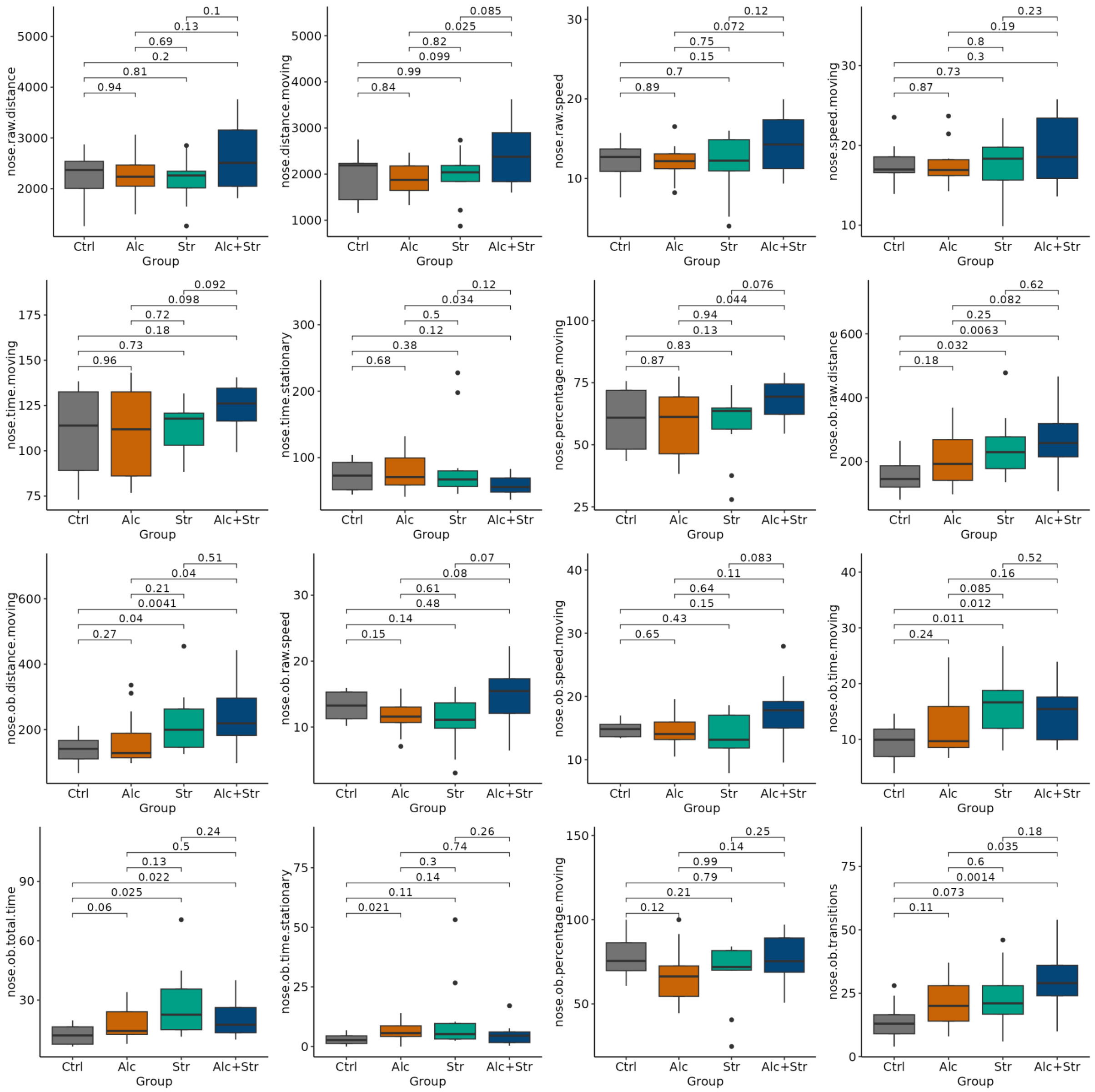
0.1
0.13
0.69
0.2
0.81
0.94
5000
4000
3000
2000
nose.raw.distance
Ctrl
Alc
Str
Alc+Str
Group
0.085
0.025
0.82
0.099
0.99
0.84
5000
4000
3000
2000
1000
nose.distance.moving
Ctrl
Alc
Str
Alc+Str
Group
0.12
0.072
0.75
0.15
0.7
0.89
30
20
10
nose.raw.speed
Ctrl
Alc
Str
Alc+Str
Group
0.23
0.19
0.8
0.3
0.73
0.87
30
20
10
nose.speed.moving
Ctrl
Alc
Str
Alc+Str
Group
0.092
0.098
0.72
0.18
0.73
0.96
175
150
125
100
75
nose.time.moving
Ctrl
Alc
Str
Alc+Str
Group
0.12
0.034
0.5
0.12
0.38
0.68
300
200
100
nose.time.stationary
Ctrl
Alc
Str
Alc+Str
Group
0.076
0.044
0.94
0.13
0.83
0.87
100
75
50
25
nose.percentage.moving
Ctrl
Alc
Str
Alc+Str
Group
0.62
0.082
0.25
0.0063
0.032
0.18
600
400
200
nose.ob.raw.distance
Ctrl
Alc
Str
Alc+Str
Group
0.51
0.04
0.21
0.0041
0.04
0.27
600
400
200
nose.ob.distance.moving
Ctrl
Alc
Str
Alc+Str
Group
0.07
0.08
0.61
0.48
0.14
0.15
30
20
10
nose.ob.raw.speed
Ctrl
Alc
Str
Alc+Str
Group
0.083
0.11
0.64
0.15
0.43
0.65
40
30
20
10
nose.ob.speed.moving
Ctrl
Alc
Str
Alc+Str
Group
0.52
0.16
0.085
0.012
0.011
0.24
40
30
20
10
nose.ob.time.moving
Ctrl
Alc
Str
Alc+Str
Group
0.24
0.5
0.13
0.022
0.025
0.06
90
60
30
nose.ob.total.time
Ctrl
Alc
Str
Alc+Str
Group
0.26
0.74
0.3
0.14
0.11
0.021
75
50
25
0
nose.ob.time.stationary
Ctrl
Alc
Str
Alc+Str
Group
0.25
0.14
0.99
0.79
0.21
0.12
150
100
50
nose.ob.percentage.moving
Ctrl
Alc
Str
Alc+Str
Group
0.18
0.035
0.6
0.0014
0.073
0.11
75
50
25
0
nose.ob.transitions
Ctrl
Alc
Str
Alc+Str
Group

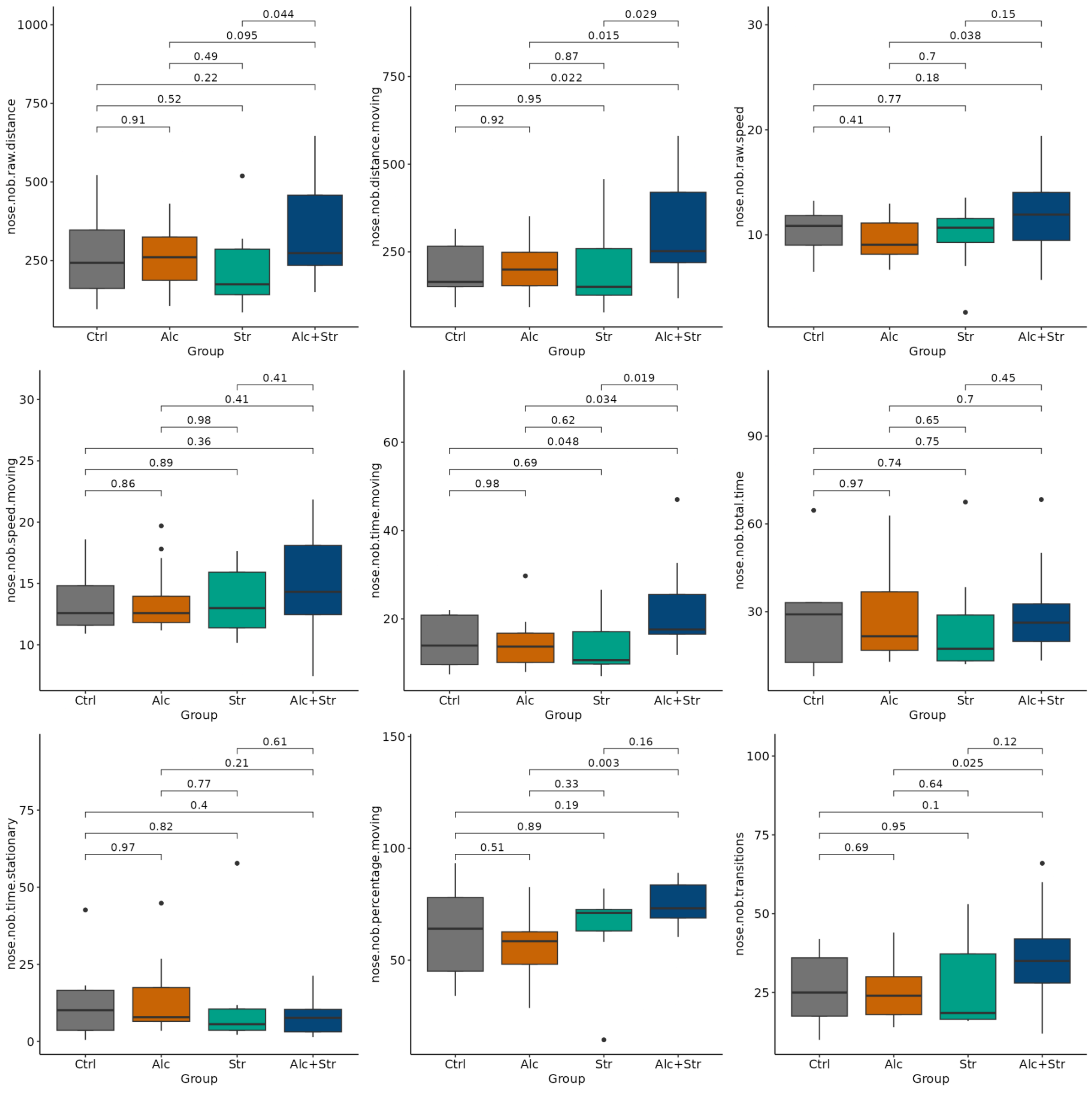

nose.nob.raw.distance
0.044
0.095
0.49
0.22
0.52
0.91
Group
Ctrl
Alc
Str
Alc+Str
nose.nob.distance.moving
0.029
0.015
0.87
0.022
0.95
0.92
nose.nob.raw.speed
0.15
0.038
0.7
0.18
0.77
0.41
nose.nob.speed.moving
0.41
0.41
0.98
0.36
0.89
0.86
nose.nob.time.moving
0.019
0.034
0.62
0.048
0.69
0.98
nose.nob.total.time
0.45
0.7
0.65
0.75
0.74
0.97
nose.nob.time.stationary
0.61
0.21
0.77
0.4
0.82
0.97
nose.nob.percentage.moving
0.16
0.003
0.33
0.19
0.89
0.51
nose.nob.transitions
0.12
0.025
0.64
0.1
0.95
0.69

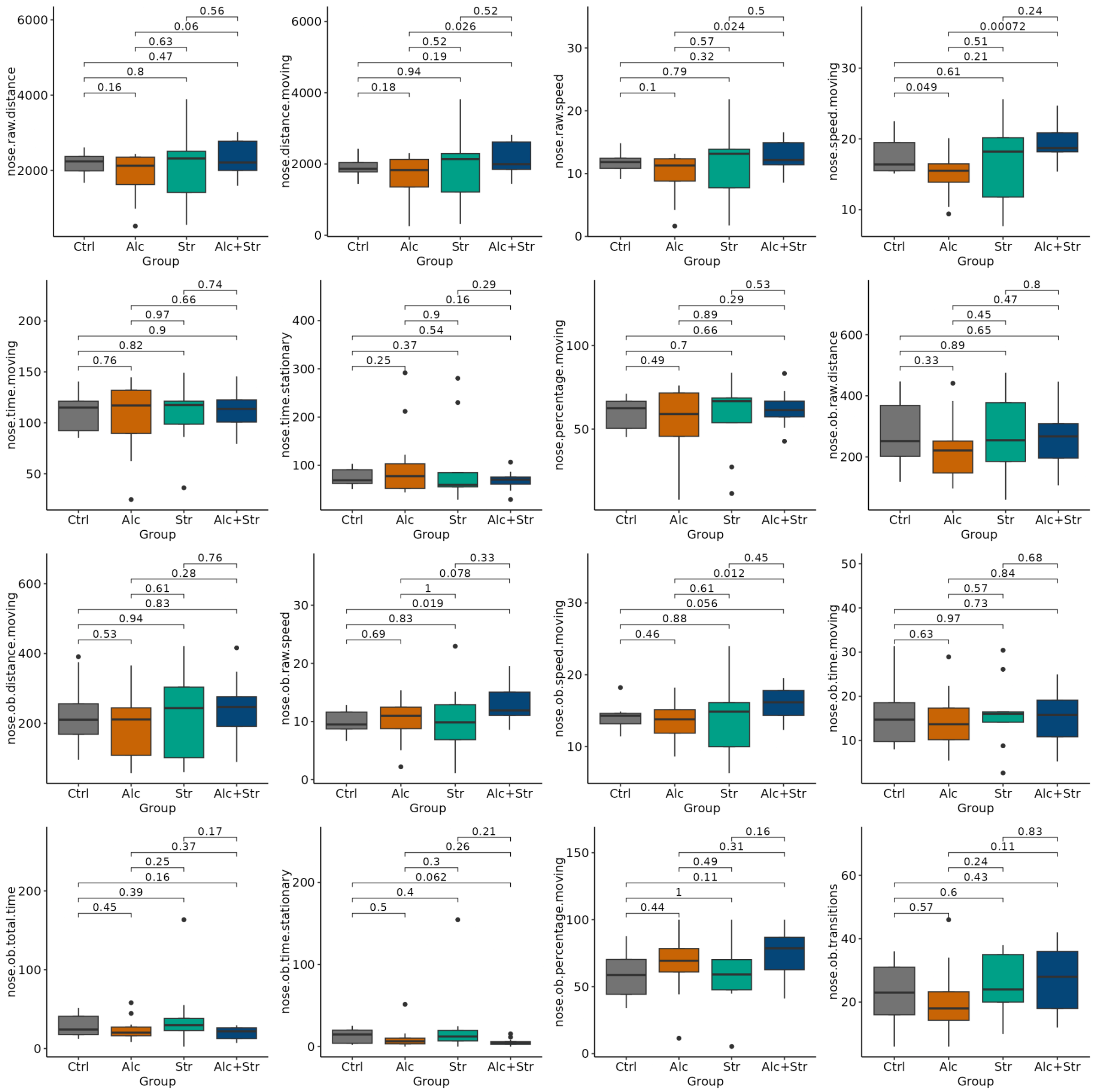
nose.raw.distance
6000
4000
2000
0.56
0.06
0.63
0.47
0.8
0.16
Ctrl
Alc
Str
Alc+Str
Group
nose.distance.moving
6000
4000
2000
0
0.52
0.026
0.52
0.19
0.94
0.18
Ctrl
Alc
Str
Alc+Str
Group
nose.raw.speed
30
20
10
0
0.5
0.024
0.57
0.32
0.79
0.1
Ctrl
Alc
Str
Alc+Str
Group
nose.speed.moving
30
20
10
0.24
0.00072
0.51
0.21
0.61
0.049
Ctrl
Alc
Str
Alc+Str
Group
nose.time.moving
200
150
100
50
0.74
0.66
0.97
0.9
0.82
0.76
Ctrl
Alc
Str
Alc+Str
Group
nose.time.stationary
400
300
200
100
0.29
0.16
0.9
0.54
0.37
0.25
Ctrl
Alc
Str
Alc+Str
Group
nose.percentage.moving
100
50
0.53
0.29
0.89
0.66
0.7
0.49
Ctrl
Alc
Str
Alc+Str
Group
nose.ob.raw.distance
600
400
200
0.8
0.47
0.45
0.65
0.89
0.33
Ctrl
Alc
Str
Alc+Str
Group
nose.ob.distance.moving
600
400
200
0.76
0.28
0.61
0.83
0.94
0.53
Ctrl
Alc
Str
Alc+Str
Group
nose.ob.raw.speed
30
20
10
0
0.33
0.078
1
0.019
0.83
0.69
Ctrl
Alc
Str
Alc+Str
Group
nose.ob.speed.moving
30
20
10
0.45
0.012
0.61
0.056
0.88
0.46
Ctrl
Alc
Str
Alc+Str
Group
nose.ob.time.moving
50
40
30
20
10
0.68
0.84
0.57
0.73
0.97
0.63
Ctrl
Alc
Str
Alc+Str
Group
nose.ob.total.time
200
100
0
0.17
0.37
0.25
0.16
0.39
0.45
Ctrl
Alc
Str
Alc+Str
Group
nose.ob.time.stationary
200
100
0
0.21
0.26
0.3
0.062
0.4
0.5
Ctrl
Alc
Str
Alc+Str
Group
nose.ob.percentage.moving
150
100
50
0
0.16
0.31
0.49
0.11
1
0.44
Ctrl
Alc
Str
Alc+Str
Group
nose.ob.transitions
60
40
20
0.83
0.11
0.24
0.43
0.6
0.57
Ctrl
Alc
Str
Alc+Str
Group

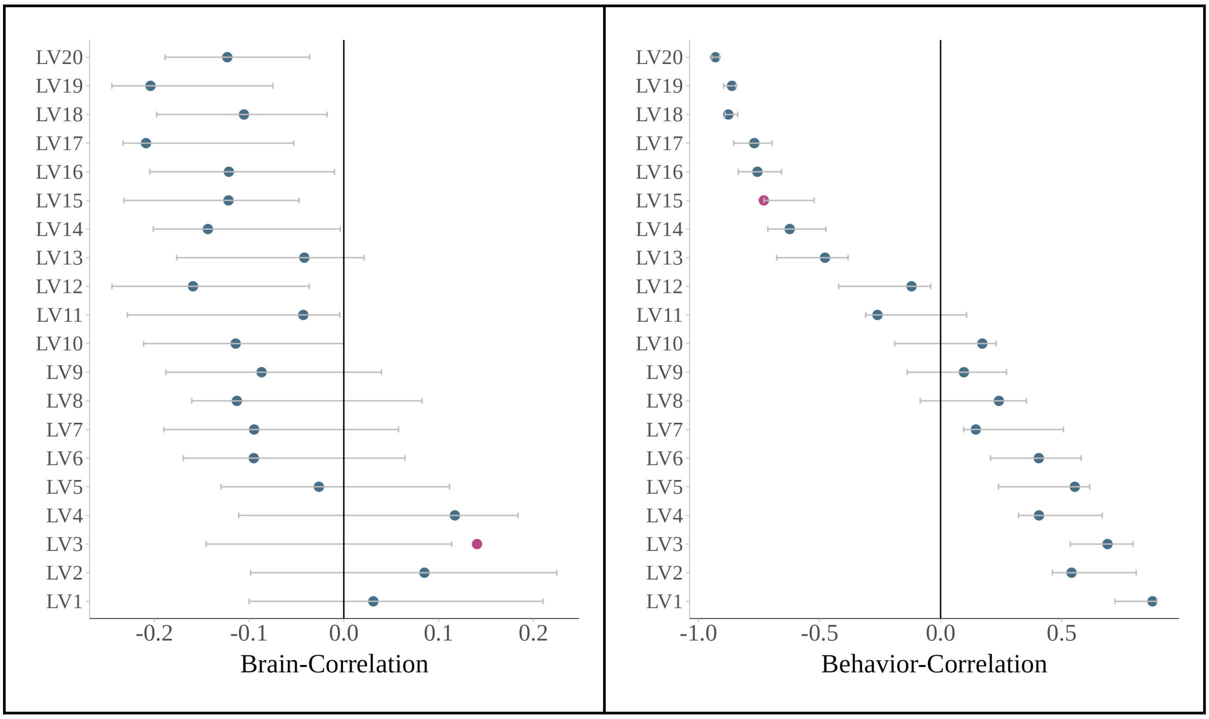

LV20
LV19
LV18
LV17
LV16
LV15
LV14
LV13
LV12
LV11
LV10
LV9
LV8
LV7
LV6
LV5
LV4
LV3
LV2
LV1
-0.2
-0.1
0.0
0.1
0.2
Brain-Correlation
-1.0
-0.5
0.0
0.5
Behavior-Correlation
