## Supplementary discussion for "The effect of chronic stress and chronic alcohol intake on behavior, brain structure, and functional connectivity in a rat model"

The conditioned place preference (CPP) task revealed that all groups exhibited an equal preference for the ethanol-paired and water-paired compartments. This contradicts findings from studies that report stress-induced ethanol preference in humans and rodent models, but is consistent with research showing age-dependent effects, where they have found that adolescent rats show no ethanol preference (1–4). Stress-induced ethanol preference could be related to other factors, such as individual variations in stress response and compensatory ethanol intake, as well as social and environmental factors that may be unique to humans (5,6). In this sense, classification models can help to longitudinally identify subgroups with distinct intake patterns that deviate from the overall group mean, as suggested by previous results in our laboratory (7).

The findings from the Elevated Plus Maze (EPM) task suggest that ethanol intake and chronic restraint stress (CRS) have differential effects on anxiety-like behaviors in male and female rats. Notably, while the overall anxiety index did not vary significantly among groups, detailed analysis revealed that female rats exposed to ethanol (EtOH+/CRS- and EtOH+/CRS+) exhibited reduced movement and speed, and an increased stationary and total time spent but only in one closed arm. Meanwhile, males exposed to both ethanol and CRS exhibit a more complex pattern with increased exploration in the center and open arms but also increased stationary behavior in the closed arms. These results might imply that ethanol, even after a long withdrawal period, reduces the natural exploratory behavior in females (8,9), while has mixed effects in males depending on their exposure to chronic stress.

Similarly, the results from the Novel Object Recognition (NOR) task showed a reduced discrimination ratio in the male EtOH-/CRS+ group, indicating impaired recognition and memory performance under conditions of chronic stress (10–13). This observation aligns with previous research demonstrating that chronic stress can adversely affect hippocampal function, leading to memory deficits that vary between sexes (13,14). Notably, male rats exposed to both ethanol and stress (EtOH+/CRS+) exhibited increased locomotor activity, characterized by greater speed and distance traveled. This behavior may suggest hyperactivity or enhanced exploratory tendencies (15), consistent with studies investigating the effects of chronic stress alone (16) and those examining ethanol exposure independently (17).

Our results further suggest that while the combination of chronic ethanol intake and chronic stress exacerbated the effects on local brain anatomy and functional connectivity network, we also found that each intervention showed distinct patterns of structural and functional neuroadaptations, highly affected by chronic stress more than ethanol intake. In the structural analysis, when we compared the effect of both interventions against the groups with only one intervention (chronic stress or ethanol intake), we found that chronic stress produced an increased volume in Cg2 and RSGc along with a decrease of ventral hippocampus and amygdala, as well as more extensive alterations in local volume within hippocampus, caudate-putamen and amygdala, which are essential neuroanatomical substrates of the stress-response and drug-seeking systems, and intimately involved in processes such as learning, memory, reward and anxiety-like behaviors, which also usually involve more cortical and limbic regions.(18–21) Moreover, we found altered functional connectivity between cortical and subcortical regions (e.g., amygdala-thalamus, amygdala-orbitofrontal cortex, hippocampus-cerebellum, caudate-putamen-orbitofrontal) in the chronic stress group, which is related to both reward systems in addiction and chronic stress. These changes have been previously noted(20), mostly with an FC-decreased pattern, similar to what we observed. However, our findings further reveal that these effects are sex-dependent.(22,23) Therefore, even though chronic stress did not seem to have an effect on ethanol intake, it may be affecting the addiction processes by either influencing dependency or relapse. This could give us an insight into the addictive effect of pathological chronic stress on morphology, functionality, and memory that might be caused by microstructural processes.(6,24)).

### Limitations

Limitations encompass the technical problems with the MRI scanner which resulted in the loss of seven images for the T2 and T3 points. However, this did not affect the overall results, as mixed model effects were used to account for missing data. Additionally, the limited number of acquisition points and the absence of a post-withdrawal time point may have led to an over-interpretation of the findings. While measuring blood corticosterone levels was valuable for validating the stress model, taking these measurements at multiple time points throughout the study would have provided a more comprehensive understanding of hormonal changes.
